# Real-time identification of carbapenemase-producing *Klebsiella pneumoniae* lineages and outbreak detection using FT-IR ATR

**DOI:** 10.1101/2024.08.12.607659

**Authors:** Ana Beatriz Gonçalves, Valquíria Alves, Isabel Neves, Maria Antónia Read, Natália Pinheiro, Anna Emilie Henius, Henrik Hasman, Luísa Peixe, Ângela Novais

## Abstract

Expansion of carbapenemase-producing *Klebsiella pneumoniae* (CP-*Kp*) is driven by nosocomial dissemination, and effective infection control depends on timely and reliable typing data. Here, we evaluated our previously developed Fourier-transform infrared spectroscopy (FT-IR) with attenuated total reflectance (ATR) workflow for real-time typing of *Kp* capsular (KL)-types and lineages to support infection control. FT-IR spectra were acquired from Columbia agar with 5% sheep blood cultures of all CP-*Kp* infection isolates (n=136) from hospitalized patients at a northern Portugal hospital (April 2022 – March 2023), and analyzed using automated machine-learning (ML) classification models. Typing results were confirmed by *wzi* sequencing, MLST and/or WGS. FT-IR typing on Columbia agar plates showed 73% sensitivity, 79% specificity and 74% accuracy. Our method correctly typed 94% of typeable isolates (78/83), from which 87% were comunicated in <24h. Sixty percent of non-typeable isolates were considered false negatives, but the majority (66%) was correctly predicted when re-tested in Mueller-Hinton agar, improving sensitivity (92%), specificity (76%) and accuracy (89%) of *Kp* typing. Three *Kp* lineages (ST147-KL64, ST15-KL19, ST268-KL20) represented 74% of the sample, with ST268-KL20 causing an outbreak in Neonatal Intensive Care unit, quickly recognized by FT-IR enabling immediate infection control measures. Epidemiological links between patiens were mostly found on medical, surgical and urology units, using EpiLinx software. Most isolates (98%) produced KPC-3. Our FT-IR ATR ML-based typing workflow demonstrated high performance standards in real-time and high adaptability to clonal dynamics. The unprecedent time-to-response (same day of species identification) represents an opportunity to implement timely and effective infection control measures.

**Importance:** This study represents the first prospective and real-time evaluation of FT-IR spectroscopy to type multidrug resistant *Klebsiella pneumoniae* to support surveillance and infection control. We demonstrate a high sensitivity, specificity and accuracy of a previously developed workflow that allows precise identification of *K. pneumoniae* lineages. The adaptability to changes in clonal dynamics and bacterial typing in <24h offer significant advantages in both high- and low-income countries for a timely infection control and improvement of antimicrobial resistance management.

## Introduction

*Klebsiella pneumoniae* (*Kp*) is a menace to most healthcare institutions worldwide because of the increasing incidence of multidrug resistance (MDR) phenotypes and, particularly, carbapenemase-producing (CP) isolates resulting in difficult-to-treat infections in hospitalized patients (1, 2). Previous studies have shown that the expansion of CP-*Kp* is essentially driven by nosocomial dissemination caused, in part, by ineffective measures of infection control (3, 4). Despite the stringent guidelines aimed at controlling the dissemination of carbapenemase producers (5), there is a large discrepancy between institutions and countries in their implementation due to variations in technical expertise, economic resources and human capacity (6, 7). Besides, early detection of transmission chains is key to control putative nosocomial outbreaks and prevent health complications, but the absence or the long time-to-response of CP-*Kp* strain typing methods to support timely infection control decisions (7, 8) contribute to the current scenario.

Whole genome sequencing (WGS) is the gold-standard method for strain typing and outbreak detection (8–10). Despite the potential cost-benefit of integrating WGS in the clinical practice, this technology remains challenging for institutions with low economic resources and time-to-result can be as low as 2-3 days (11) but more often takes one week or more to provide interpreted reports, which is suboptimal for real-time control (12, 13). Nanopore sequencing might well decrease time-to-response of WGS but still requires wet and dry lab expertise (14, 15).

Fourier-transformed infrared spectroscopy (FT-IR) is known to support strain typing of several clinically relevant species, including *Kp* (16, 17). The simplicity of the procedure, the quick time-to-response (including the same day than bacterial identification) and the low cost (inexpensive equipment requiring low maintenance, and few consumables needed for the procedure) are attractive features for routine implementation (18–20). Most available studies on FT-IR typing and application to outbreak detection and epidemiological surveillance of CP-*Kp* are retrospective (21, 22), while prospective studies in real-time contexts are still missing. We have recently developed a simple (similar to that of matrix-assisted desorption ionization time-of-flight mass spectrometry [MALDI-TOF MS]), fast (typing result in <10 minutes), reproducible and automated workflow of analysis using FT-IR with attenuated total reflection (ATR), based on a machine-learning (ML) classification model able to discriminate and identify up to 36 *Kp* KL-types of MDR *Kp* globally spread clonal lineages (19, 20). Here, we aim to evaluate the performance of our previously developed FT-IR workflow for typing of *Kp* KL-types and lineages and combine the results with epidemiological analysis to support infection control in real-time.

## Materials and Methods

### Study design

We established a partnership with a 370-bed district hospital located in the northern region of Portugal, serving a population of 317,910 inhabitants from the municipalities of Matosinhos, Póvoa do Varzim, and Vila do Conde. This hospital provides healthcare services across various specialties, including medical, surgical, emergency (adult and pediatric), and outpatient departments. The hospital’s Infection Control and Antimicrobial Resistance Department implements policies and interventions to minimize the spread of antimicrobial resistance. In 2023, the hospital reported a 30.1% prevalence of carbapenemase-producing *Enterobacteriaceae* (CPE) in invasive strains, a 10.5% CPE colonization rate, and an average of 40 patients per day isolated due to CPE infection or colonization.

For the purpose of this study, we focused on assessing within-hospital transmission chains of CP-*Kp*. For that, we included all (n=136) CP-*Kp* identified in infection sites from 127 hospitalized patients during a one-year period (April 2022 – March 2023). Additional isolates (n=4 from rectal colonization) were considered in the context of outbreak investigation.

To avoid extra costs and time/personnel dedicated, we implemented our FT-IR workflow as a service, whereby all CP-*Kp* were identified and sent to our lab by a transport service that assured delivery between 11h00 AM and 16h00 PM up to three times/week. All isolates were immediately registered and processed, and the results were sent to the hospital at the earliest convenience through a report sent by email to the Directors of the Microbiology and Infection Control Unit departments. All relevant epidemiological information was collected and shared for integrated epidemiological and outbreak analysis (see details below).

### Identification of CP-*Kp* isolates

According to routine procedures, sputum (15/136; 11%), pus (15/136; 11%), blood cultures (11/136; 8%) and other products (17/136; 13%) such as ocular exudate, ascitic fluid, bone/tissue biopsy, and pleural fluid human specimens were plated in Columbia agar with 5% sheep blood (bioMérieux, France) while urine samples (78/136; 57%) were plated in Cystine-lactose-electrolyte-deficient (CLED) agar (Frilabo, Portugal). Species identification was performed by MALDI-TOF MS (VITEK MS, bioMeriéux, France) and/or by VITEK 2 (bioMeriéux, France), together with antimicrobial susceptibility profiles. Screening for carbapenemase production was performed by a multiplex immunochromatographic assay (O.K.N.V.I. RESIST-5, Coris BioConcept, Belgium). All CP-*Kp* from Columbia agar with 5% sheep blood plates were sent directly to our research lab, whereas those from CLED agar were subcultured in Columbia agar with 5% sheep blood prior to shipment.

### FT-IR spectra acquisition and analysis

At the day of arrival, FT-IR ATR spectra were acquired using a Spectrum Two instrument (Perkin-Elmer, USA) and a previously described workflow (20). In brief, a single isolated colony picked from recently cultured (<24h) Columbia agar with 5% sheep blood plates received from hospital was applied in the ATR crystal of the equipment as a thin uniform layer, air-dried for 3 to 5 minutes, and three technical replicates were acquired (<5 min per isolate). Spectra were preprocessed and analysed by a previously developed and validated automated analysis workflow integrated in the Clover MS Data Analysis software (Clover Bioanalytical Software, Spain) (20). Using this approach, spectra were compared with those from a spectral database representing main MDR *Kp* lineages causing human infections. We use supervised ML classification models that are trained using a known and well-characterized training dataset, and subsequently tested with new samples from a validation dataset. These models were developed to enable classification and identification of *Kp* MDR high-risk lineages by an accurate discrimination of capsular (KL)-types (20).

### Predictive models used in this study

Two different classification models were used. We used our previously developed random forest (RF) classification model (Model 1) that can differentiate and identify 33 KL-types of globally disseminated MDR *Kp* with a high accuracy (89%), sensitivity (88%) and specificity (92%) (20). This model was updated during the study period to improve robustness (by increasing the number of spectra per class) and to increase coverage (by including additional KL-types). This allowed prediction of new KL-types circulating in this hospital. The updated RF classification model (Model 2) included 302 isolates (1815 spectra) and was designed to allow differentiation of 25 KL-types, representing worldwide spread and locally circulating KL-types. This new model was validated with spectra from a *validation set* comprising 141 well characterized isolates from Portugal and Spain, as previously described (see Supplementary Material) (20).

### Validation of FT-IR typing results

Results from the RF classification models were given by a prediction score for each KL-type. According to the criteria previously established (20), an isolate was considered typeable when the probability score of the first predicted KL-type (P1) is >25% and the difference between P1 and the probability score of the second predicted KL-type (P2) is ≥10%. When these conditions were not met, the isolate was considered non-typeable by the model, i.e., it had a KL-type different from those of the RF model. Non-typeable isolates were re-isolated in Mueller-Hinton (MH) agar (bioMérieux, France; HiMedia Laboratories, Germany) in standardized conditions (37°C for 18h), and FT-IR spectra acquisition and typing were repeated.

Sequencing of the *wzi* gene was used as the reference method to infer KL-type according to the BIGSdb Pasteur database (https://bigsdb.pasteur.fr/) (23). Multi-locus sequence typing (MLST) was performed in representative isolates of each KL-type to confirm correlation with major known *Kp* lineages. Clonal relatedness of the predominant lineages was confirmed by pulsed-field gel electrophoresis (*Xba*I-PFGE), as previously described (24).

Typeable isolates’ results were communicated to the hospital within 45min to 24h, except for cases with lower prediction scores (P1=25-27%) and/or infrequent KL-types. These isolates and all the non-typeable were only communicated after confirmation by the reference methods (48-72h).

### Evaluation of FT-IR typing performance

We evaluated the performance of FT-IR workflow for real-time typing by calculating the sensitivity, specifity, and accuracy for KL-type identification, as described (20). For this purpose, we considered 131 out of the 136 isolates. For the remaining five isolates there were unexpected delays in spectra acquisition and the workflow could not be applied as predicted. Prediction results were considered as true positives (TP, when FT-IR typeable with a result concordant with *wzi* sequencing), true negatives (TN, when non-typeable by FT-IR with an inferred KL-type was not present in the model), false positives (FP, when typeable by FT-IR with an inferred KL-type different from the result obtained) and false negatives (FN, when non-typeable by FT-IR with an inferred KL-type included in the model). Sensitivity, specificity, and accuracy were evaluated (20).

### Epidemiological analysis and outbreak investigation

Putative transmission links between all patients with infections caused by the predominant *Kp* lineages (n=95) and related patients (n=4) in outbreak investigations were identified by analysis of epidemiological data. We considered dates of admission and discharge, date of infection, patient’s trajectory within the hospital since admission until discharge, hospitalization periods in the previous four months, and positive rectal colonization by CPE at admission. Putative epidemiological links were identified using EpiLinx, a software developed by Henius *et al*. at Statens Serum Institut from Copenhagen, Denmark (available at https://github.com/AnnaMille/EpiLinx/) (25). A putative direct link was established when patients infected by the same *Kp* lineage were hospitalized in the same unit at the same time. Indirect links were inferred when patients infected by the same *Kp* lineage were hospitalized in the same unit but not at the same time (14 days difference or less).

Outbreak investigation included rectal colonization screening for CPE of all related patients and environmental screening of neonatal intensive care unit (NICU). Samples from one incubator (one swab after cleaning process), three mattresses (four swabs from the surface and three samples of foam), and two sinks (five swabs each) were collected and screened for the CP-*Kp*. Samples were enriched in Buffered Peptone Water (Liofilchem, Italy) for 1h at room temperature, followed by enrichment for 24h at 37°C and further plated in HiCrome UTI Agar (HiMedia Laboratories, Germany)/Simmons Citrate agar (Liofilchem, Italy) + *myo*-inositol (Thermo Fisher Scientific, USA) supplemented or not with imipenem (1 mg/L) (Merck, Germany).

### Whole genome sequencing

WGS was performed in 16 isolates: eight representing different KL-types, to confirm capsular (whole *cps*) – biochemical (FT-IR) correlations, and resistance/virulence genotypic profiles; two isolates involved in the NICU outbreak and six non-outbreak isolates from the same KL-type (one per month since the beginning of the study) to confirm genetic relatedness.

Genomic DNA was extracted using Wizard Genomic DNA purification kit (Promega Corporation, Madison, WI) and the final concentration was assessed with a Qubit 3.0 Fluorometer (Invitrogen, Thermo Fisher Scientific, USA). Genomic DNA was then sequenced with Genome Sequencer Illumina NovaSeq (Illumina, San Diego, CA, USA), using NovaSeq 6000 S4 PE150 XP mode, at Eurofins Genomics (https://eurofinsgenomics.eu/). Raw sequence data quality was confirmed with FastQC v0.12.1 (26), using the default parameters. SPAdes v3.13.0 was used for assembly, and QUAST v5.0.2 for assembly quality assessement, at Bacterial and Viral Bioinformatics Resource Center (BV-BRC, https://www.bv-brc.org/). Genome assemblies were analysed using Pathogenwatch v22.1.3 (https://pathogen.watch/), supported by Kaptive for capsular polysaccharide (K) and lipopolysaccharide (O) locus types and serotypes (27), BIGSdb-Pasteur (https://bigsdb.pasteur.fr/klebsiella/) for MLST, core genome (cg) MLST and cgLIN code (28, 29), Kleborate for antibiotic resistance and virulence determinants (30), and PlasmidFinder from the Centre for Genomic and Epidemiology for plasmid replicon typing (31).

Our outbreak-related genomes were compared to genomes from Pathogenwatch public database, using cgMLST single linkage clustering, and the most closely related (17 genomes with <10 allele differences within the 629 genes included in cgMLST scheme) were selected for phylogenetic and epidemiological analysis. Neighbour-joining tree based in the pairwise single nucleotide polymorphism (SNP) distances between genomes based on a concatenated alignment of 1,972 core genes was generated, using Pathogenwatch (32).

## Results

### FT-IR typing performance in real-time

Our typing workflow based only on Columbia agar with 5% sheep blood cultures showed 73% of sensitivity, 79% of specificity and 74% of accuracy to identify *Kp* KL-types. Figure 1 represents typeability and time-to-response obtained in this study. Out of the 131 isolates evaluated, 83 (63%) were typeable and 48 (37%) were non-typeable by the model used. Most typeable isolates were typed correctly (n=78/83, 94%), whereas 5 (6%) were considered FP results based on reference methods. The result was communicated in <24h in 87% of the cases, where 43% of these on the day of arrival, and only two resulted in incorrect predictions. The remaining isolates (n=11/83, 13%) were communicated in 48-72h after confirmation by reference methods. These belonged to infrequent KL-types (e.g. KL105 and KL23) or yielded lower prediction scores (P1=25-27%), and only three were FP results.

**Figure 1.**
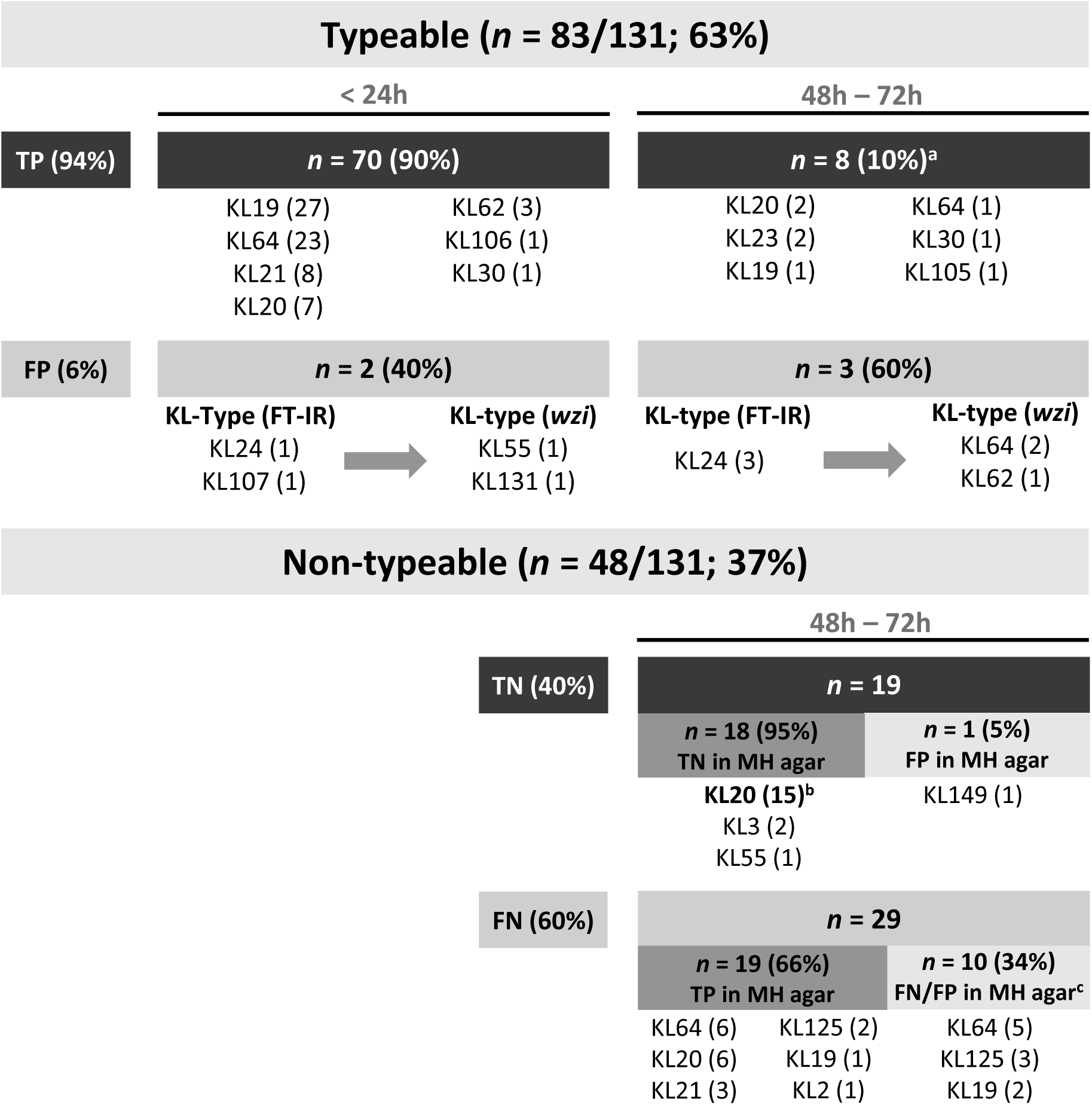
Typeability and time-to-response of CP-*Kp* FT-IR typing workflow in real-time. Isolates are considered typeable when probability scores match the defined criteria (P1>25% and P1-P2>10%) and non-typeable when these criteria are not met. TP = true positive, FP = false positive, TN = true negative, and FN = false negative. ^a^Isolates with FT-IR typing results with P1=25-27% and/or infrequent KL-types. ^b^Isolates typed with RF Model 1. ^c^Four KL64 and two KL19 isolates were non-mucoid variants. Only one isolate (KL125) was a FP in MH agar.

Within the non-typeable isolates, 19 (40%) corresponded to four KL-types that were not included in Model 1, and were thus considered TN results. Most of these were KL20 isolates (15/19; 79%) that were recognized as highly related by FT-IR spectroscopy, thus considered putative outbreak strains (Supplementary Figure 1). FN results (60%) included six KL20 isolates that were not identified by Model 2, and a few non-mucoid variants of KL64 and KL19 isolates. Interestingly, 19 (66%) FN isolates were correctly predicted after a re-isolation step in MH agar.

Thus, when considering prediction results from Columbia agar with 5% sheep blood and re-isolation of non-typeable isolates in MH agar, we obtained a combined performance of 92% of sensitivity rate, 72% of specificity and 88% of accuracy, and most isolates (74%) would be correctly typed in <24h (Figure 2).

**Figure 2.**
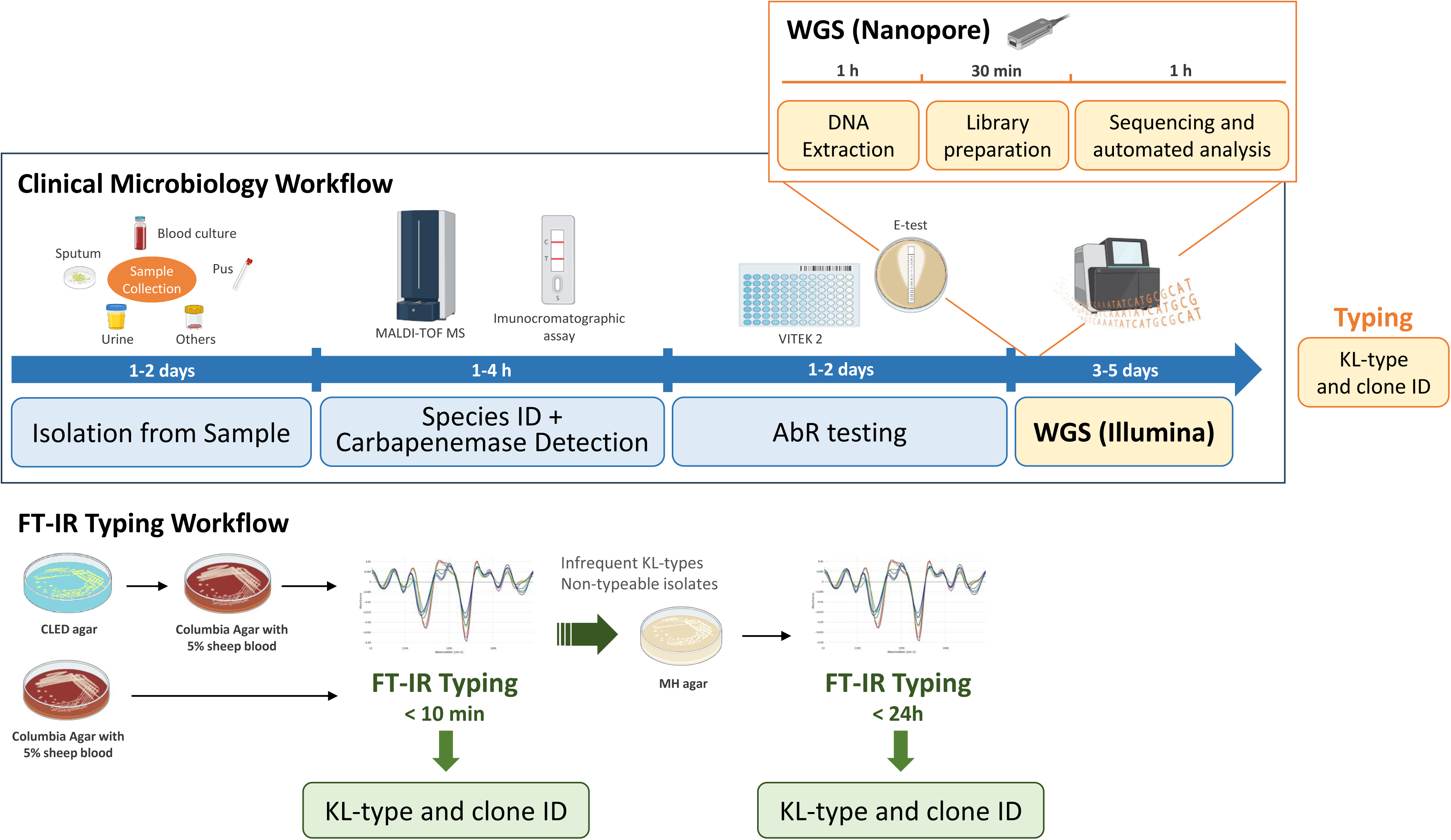
Representation of the clinical microbiology workflow timeline for pathogen identification and typing using FT-IR and WGS. Timing and time-to-response of FT-IR *vs* whole genome sequencing (both Illumina or Nanopore sequencing) are compared.

### Diversity of CP-*Kp* lineages and identification of transmission chains

Three *Kp* lineages were predominant among a total of 16, representing 74% of the sample (n=101/136): ST147-KL64 (28.7%), ST15-KL19 (23.5%), and ST268-KL20 (22.1%) (Table 1). FT-IR spectroscopy correctly identified 86% of the isolates from these predominant lineages. All isolates from each of these lineages shared highly similar PFGE profiles (2-5 band differences, Supplementary Figure 2), supporting clonal relatedness recognized by FT-IR. In addition, epidemiological links were established between patients with infections caused by these predominant *Kp* lineages, most of them related to medical (23 links) and surgical (17 links) units, and Urology (8 links) (Figure 3).

**Table 1.** CP-*Kp* clone and KL-type diversity, frequency and FT-IR typeability.

| ST | KL-type | Number of isolates (%) |
| --- | --- | --- |
| ST147 | KL64 | 39 (28.7%)<sup>a</sup> |
| ST15 | KL19 | 32 (23.5%)<sup>a</sup> |
| ST268 | KL20 | 30 (22.1%) |
| ST323 | KL21 | 11 (8.1%) |
| ST219 | KL125 | 7 (5.1%)<sup>a</sup> |
| ST45 | KL62 | 4 (2.9%) |
| ST13 | KL3 | 2 (1.5%) |
| ST280 | KL23 | 2 (1.5%) |
| ST268 | KL55 | 2 (1.5%) |
| ST25 | KL2 | 1 (0.7%) |
| ST29 | KL30 | 1 (0.7%) |
| ST2328 | KL30 | 1 (0.7%) |
| ST11 | KL105 | 1 (0.7%) |
| ST101 | KL106 | 1 (0.7%) |
| ST667 | KL149 | 1 (0.7%) |
| ST1451 | KL131 | 1 (0.7%) |
| <sup>a</sup>FT-IR spectroscopy workflow was not applied in 5 isolates (2 ST147-KL64, 2 ST219-KL125, and 1 ST15-KL19). |  |  |

**Figure 3.**
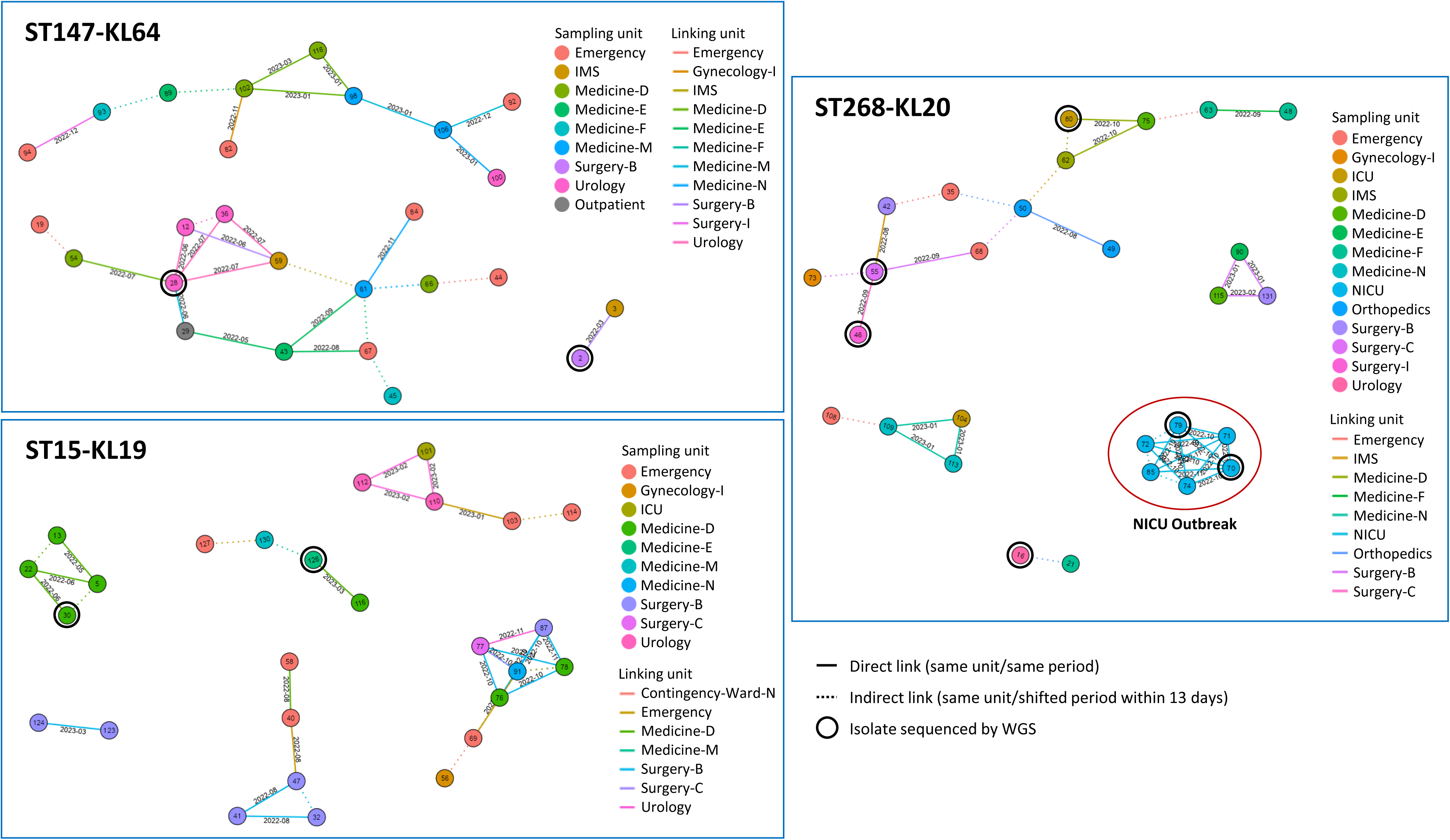
Representation of the epidemiological links between patients with infections caused by predominant CP-*Kp* lineages.

ST323-KL21, ST219-KL125 and ST45-KL62 were responsible for 8.1%, 5.1%, and 2.9% of the infections, respectively. The remaining 10% of the infections were caused by 10 infrequent lineages (n=1-2 isolates each), six of them were detected for the first time in this hospital during the course of this study.

### FT-IR supporting outbreak detection in real-time

Between October and November 2022, there was a suspicion of an outbreak in the neonatal intensive care unit (NICU). The first patient with CP-*Kp* infection was detected on October 10^th^, after which rectal colonization screenings were performed in all NICU patients yielding two positive results. At October 17^th^, two isolates from infection recovered from the same patient and two isolates from rectal colonization were sent to our lab in Columbia agar with 5% sheep blood agar plates. We confirmed immediately (<1h after reception of samples) that all isolates belonged to ST268-KL20 based on FT-IR spectral analysis. The rapid identification of the clone allowed immediate infection control measures, including transmission-based precautions, unit disinfection, clean-dirty circuits review, and environmental screening. Three additional ST268-KL20 isolates (two from rectal colonization, one from infection) were subsequently identified till November 2^nd^).

All the environmental samples collected in NICU were negative for CP-*Kp* (data not shown). The outbreak was declared resolved within 23 days. ST268-KL20 was associated with seven patients and 12 direct transmission links between NICU patients, but no epidemiological links were found between NICU patients and patients from other units. ST268-KL20 NICU strains differed in 1-5 SNPs from non-ICU strains suggesting a common non-identified source (Supplementary Table 4).

The ST268-KL20 genomes sequenced in this study had <10 allele difference within 1,972 core genes and were thus considered highly related with 17 *Kp* public genomes from human origin between 2017 and 2023 (Figure 4, Supplementary Table S4). Most of these were recovered from infection sites (82%), in the USA (88%) but also in the United Kingdom and Spain (n=1 each). Of note, one of these genomes from the USA in 2017 showed only 12-14 SNPs with our genomes. Public ST268-KL20 genomes frequently carried *bla*_KPC-3_, *bla*_NDM-1_ and/or *bla*_CTX-M-15_, or none acquired β-lactamase (n=6). All ST268-KL20 isolates had a low virulence score (0–1) and all but one presented *yersiniabactin* (*ybt* 10, ICEKp4).

**Figure 4.**
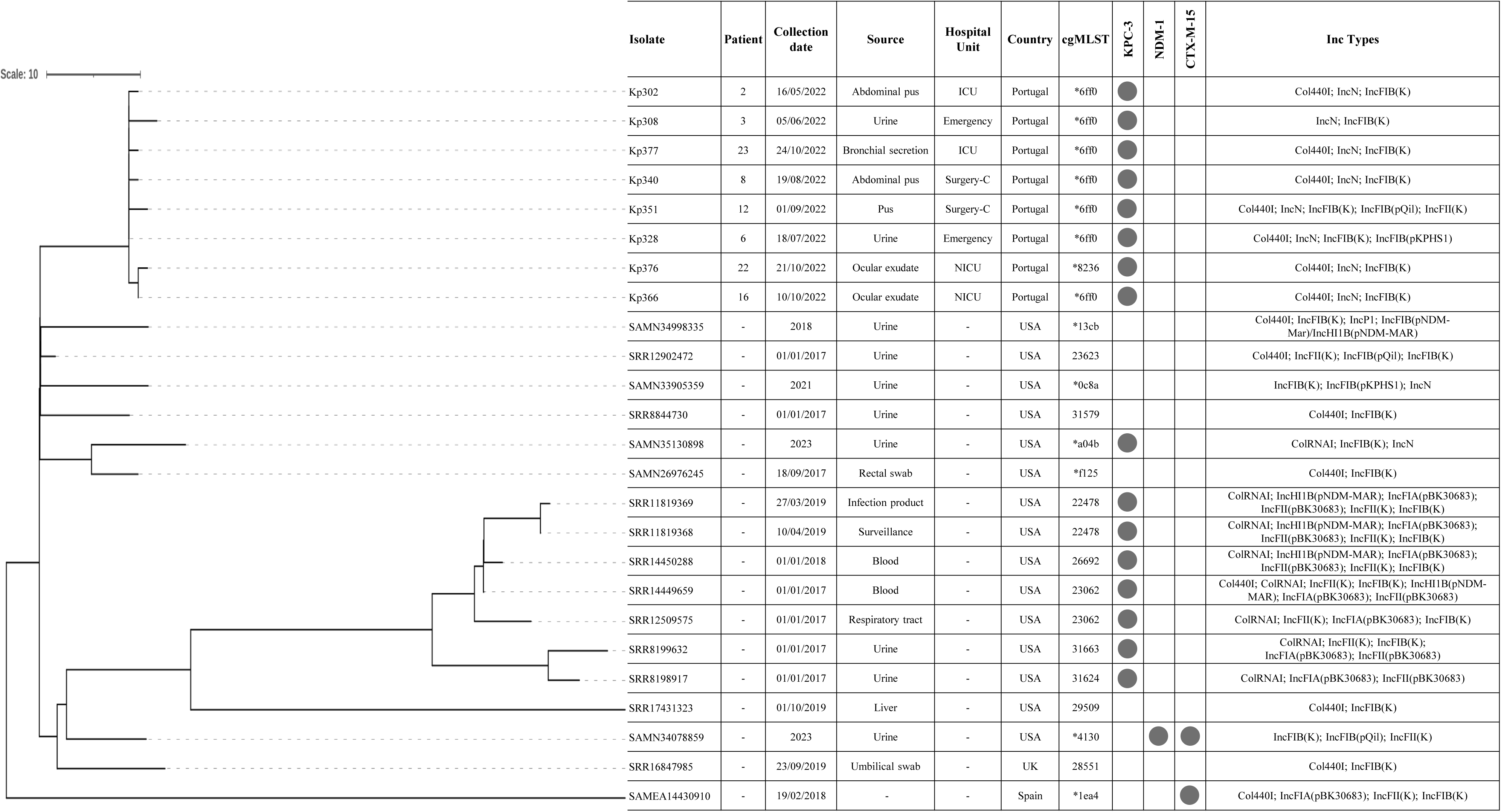
Neighbor-joining tree representing the phylogenetic relationships among the eight *Kp* ST268-KL20 from this study and 17 public genomes constructed from the Pathogenwatch pairwise-distance matrix (i.e., based on single nucleotide polymorphisms (SNPs) called in 1,972 core genes) considering >10 alleles difference. All the isolates shared the same *wzi* allele (*wzi*95), KL-type (KL20), and O-type (O2a). Pairwise SNP distances are shown in Supplementary Table 4.

### Epidemiology of resistance of *Kp* predominant clones

The vast majority of isolates produced KPC-3, whereas two isolates produced KPC-31, a KPC-3-variant, and one isolate produced IMP-22. Interestingly, the three predominant clones exhibited characteristic and distinctive antimicrobial resistance profiles to non-β-lactams, particularly to gentamicin (GN), ciprofloxacin (CIP) and trimethoprim-sulfamethoxazole (SXT). ST15-KL19 showed to be the most resistant clone, whereas ST268-KL20 was the least (Table 2). The antibiotic resistance profiles were associated with the presence of genes responsible for resistance to β-lactams (*bla*_KPC-3_, *bla*_KPC-31_, and *bla*_CTX-M-15_), GN (*strA*, *strB, aac*(*3*)*-IIa*, and *aac(6’)-Ib-cr*), CIP (*aac(6’)-Ib-cr* and *qnrS1*, together with mutations in DNA gyrase gene *gyrA* and DNA topoisomerase gene *parC*), and SXT (*dfrA14* and *sul2*) (Supplementary Figure 3).

**Table 2.**
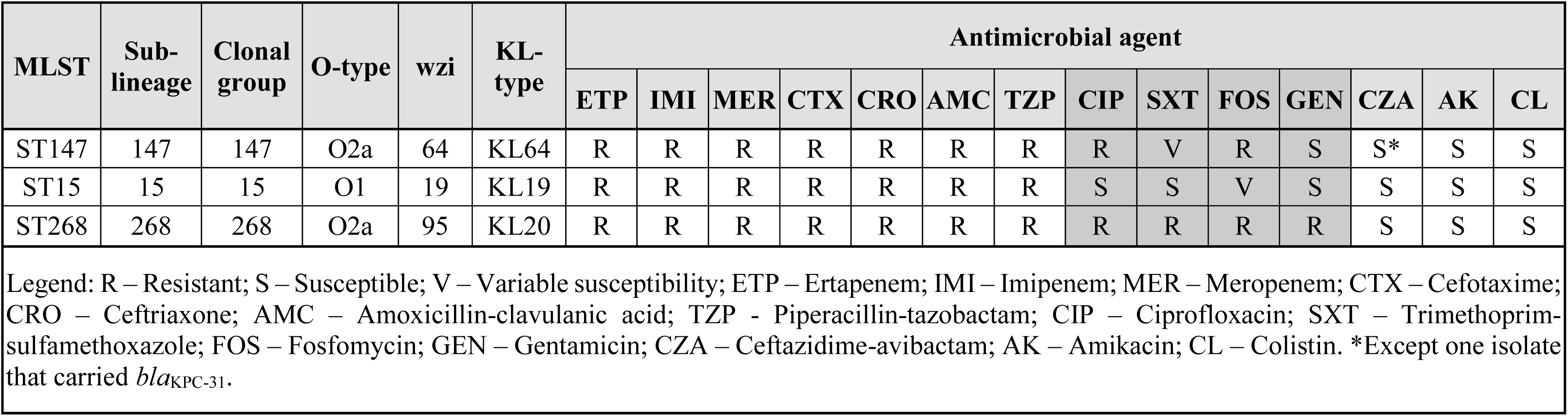
Antimicrobial resistance profiles to non-β-lactams in the three predominant CP-*Kp* lineages.

| MLST | Sub-lineage | Clonal group | O-type | wzi | KL-type | Antimicrobial agent |  |  |  |  |  |  |  |  |  |  |  |  |  |
| --- | --- | --- | --- | --- | --- | --- | --- | --- | --- | --- | --- | --- | --- | --- | --- | --- | --- | --- | --- |
|  |  |  |  |  |  | ETP | IMI | MER | CTX | CRO | AMC | TZP | CIP | SXT | FOS | GEN | CZA | AK | CL |
| ST147 | 147 | 147 | O2a | 64 | KL64 | R | R | R | R | R | R | R | R | V | R | S | S* | S | S |
| ST15 | 15 | 15 | O1 | 19 | KL19 | R | R | R | R | R | R | R | S | S | V | S | S | S | S |
| ST268 | 268 | 268 | O2a | 95 | KL20 | R | R | R | R | R | R | R | R | R | R | R | S | S | S |
| Legend: R – Resistant; S – Susceptible; V – Variable susceptibility; ETP – Ertapenem; IMI – Imipenem; MER – Meropenem; CTX – Cefotaxime; CRO – Ceftriaxone; AMC – Amoxicillin-clavulanic acid; TZP - Piperacillin-tazobactam; CIP – Ciprofloxacin; SXT – Trimethoprim-sulfamethoxazole; FOS – Fosfomycin; GEN – Gentamicin; CZA – Ceftazidime-avibactam; AK – Amikacin; CL – Colistin. *Except one isolate that carried *bla*<sub>KPC-31</sub>. |  |  |  |  |  |  |  |  |  |  |  |  |  |  |  |  |  |  |  |

## Discussion

This is the first prospective study evaluating the performance of FT-IR-based *Kp* typing in a real-time context using a routine compatible workflow. Our data demonstrates that *Kp* clonal identification based on a spectral database and a ML classification model provides not only accurate and quick results (<24h in 74% of the cases) supporting real-time surveillance, outbreak detection and infection control, but it is also flexible and adaptable to accommodate changes in the epidemiology.

The results obtained in this study and previously (20) show that FT-IR performance standards are high, and slight variations can occur depending on the nature and diversity of the population and culture conditions. The lower specificity observed in this study (79% *vs* 92%) seems to be associated with the lower number and diversity of TN of the sample (15%; most of them KL20-ST268). On the other hand, the sensitivity is affected by variations in culture conditions, as previously recognized (18, 33), as it increases from 73% to 92% when we considered results after re-isolation in MH agar. This might well happen because our spectral database is created with spectra obtained from MH agar plates (19, 20). To minimize this effect, we recommend building spectral databases and models with spectra from both Columbia agar with 5% sheep blood and MH agar cultures to better support prediction models in real-time. Furthermore, non-mucoid variants are not correctly identified since capsule is absent, which is critical for FT-IR based identification. These acapsular mutants occur by acquisition of mutations in the *cps* locus and affect a small part of the *Kp* population (6-22%), having been described for internationally important *Kp* lineages such as ST258-KL107 or ST11-KL64 (34, 35).

One of the major advantages of FT-IR over other typing techniques is the extraordinary short time-to-response. It is of highlight that we correctly identified the majority of isolates at the day of arrival (74%), which corresponds to the same day *Kp* is cultured from the clinical sample and identified by MALDI-TOF MS. A fraction of the isolates (15%) were correctly identified the day after, upon re-isolation in MH agar. This represents a still unmet time-to-response for bacterial typing that can support anticipated and more effective infection control measures. Thus, while testing in Columbia agar with 5% sheep blood agar provides maximum speed, we recommend additional re-testing in MH or vice-versa to maximize accuracy whenever prediction results are poor (P1≤27% and P1-P2<10%). This does not impact routine workflows since MH agar plates are usually available due to antibiotic susceptibility testing assays. We show that a given ML model for clonal prediction by FT-IR is plastic and adaptable to epidemiological changes and clonal evolution throughout time. In fact, we successfully retrained our previously developed model (RF Model 1) to identify ST268-KL20 during and after the outbreak, as well as other locally important clones. The new RF Model 2 yielded similar performance standards than that previously reported (20) and allowed identification of a larger number of clones of relevance for this hospital during the period of study.

Despite the great diversity among CP-*Kp* isolates identified in this hospital, three lineages (ST147-KL64, ST15-KL19 and ST268-KL20) were predominant. In fact, ST147-KL64 has been prevalent all over the country during the last decade (24, 36, 37) as one of the main vehicles of *bla*_KPC-3_ dissemination in Portugal. ST15-KL19 has been described as a minor lineage of ST15 worldwide (38), but it seems to be increasing in prevalence in the north of Portugal (Gonçalves *et al*., unpublished data). ST268-KL20 emerges in this hospital during the study period, although it has been reported in different countries, especially in Asia (39, 40). Because these lineages present distinctive antibiotic resistance profiles, their identification at an early stage is of high relevance for antimicrobial stewardship purposes. However, the lack of real-time typing tools in Portuguese hospitals is preventing the use of typing information to guide antimicrobial therapy (41).

Our molecular and epidemiological data reveal several transmission chains within the hospital, involving particularly urology, surgical and medical units. This information motivated the reinforcement of infection control measures, including: (1) review of risk assessment for hospitalized patients (specific assessment for neonates), isolation strategies (differentiated by aggressiveness of the given clone), new clean-dirty circuits definition, new rules for cleaning the environment and equipment; (2) improvements on facility’s design, layout, and infrastructure to support better hygiene and circuits; and (3) environmental screening. It is important to consider that 45% (n=43/95) of the patients with infections by one of the three predominant clones were colonized by MDR *Kp* at admission. We did not explore colonization isolates, but previous colonization is a recognized risk factor to develop infection especially if it occurs during hospitalization (42). No epidemiological links were found in 22% of the patients (n=21/95) but some patients were colonized without previous hospitalizarion record or antibiotic therapy, suggesting a role for plasmid transmission and/or frequent reintroduction from community reservoirs, as reported (41).

In conclusion, we confirm high performance standards for the application of our ML-based FT-IR typing workflow in real-time, together with a high flexibility to be adapted to local clonal dynamics. The possibility to have typing information the same day bacteria are identified from the clinical sample represents an unprecedented opportunity to guide effective infection control and antimicrobial stewardship in real-time.

## Ethics statement

This study was approved by the Comissão Local de Proteção e Segurança da informação from the Local Healthcare Unit, Matosinhos, Portugal (Refª 76/CLPSI/2022).

## Data availability Statement

EpiLinx software is available at https://github.com/AnnaMille/EpiLinx/. The sequencing data of the 16 isolates in this study have been deposited in the National Center for Biotechnology Information (https://www.ncbi.nlm.nih.gov) under the Bioproject accession number PRJNA1144909 with the accession numbers of individual isolates listed in Supplementary Figure 3 (when available during revision process).

## Suplementary material

### Validation of the RF Model 2

The RF Model 2 was challenged with a collection of 141 isolates, including 109 isolates representing 22/25 KL-types included in the model and 32 isolates of KL-types different from those included in the model (Supplementary Tables 1 and 2), and its performance was evaluated. The Model 2 exhibited 93% of sensitivity rate, 79% of specificity and 89% of accuracy in discriminating *Kp* KL-types (Supplementary Table 3).

## Supplementary Figures

**Supplementary Figure 1.**
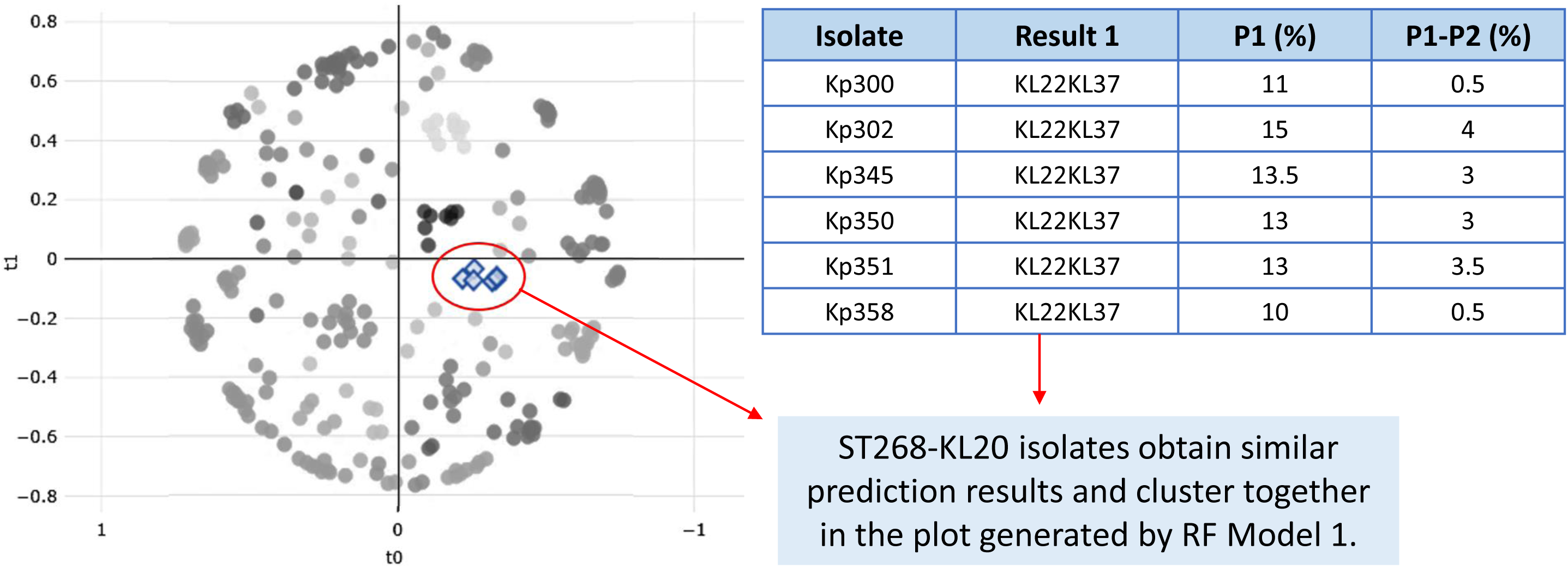
Clonal relation recognition between ST268-KL20 isolates by FT-IR spectroscopy. Spectra that support RF Model 1 are colored in shades of grey, and spectra from ST268-KL20 isolates are colored in blue.

**Supplementary Figure 2.**
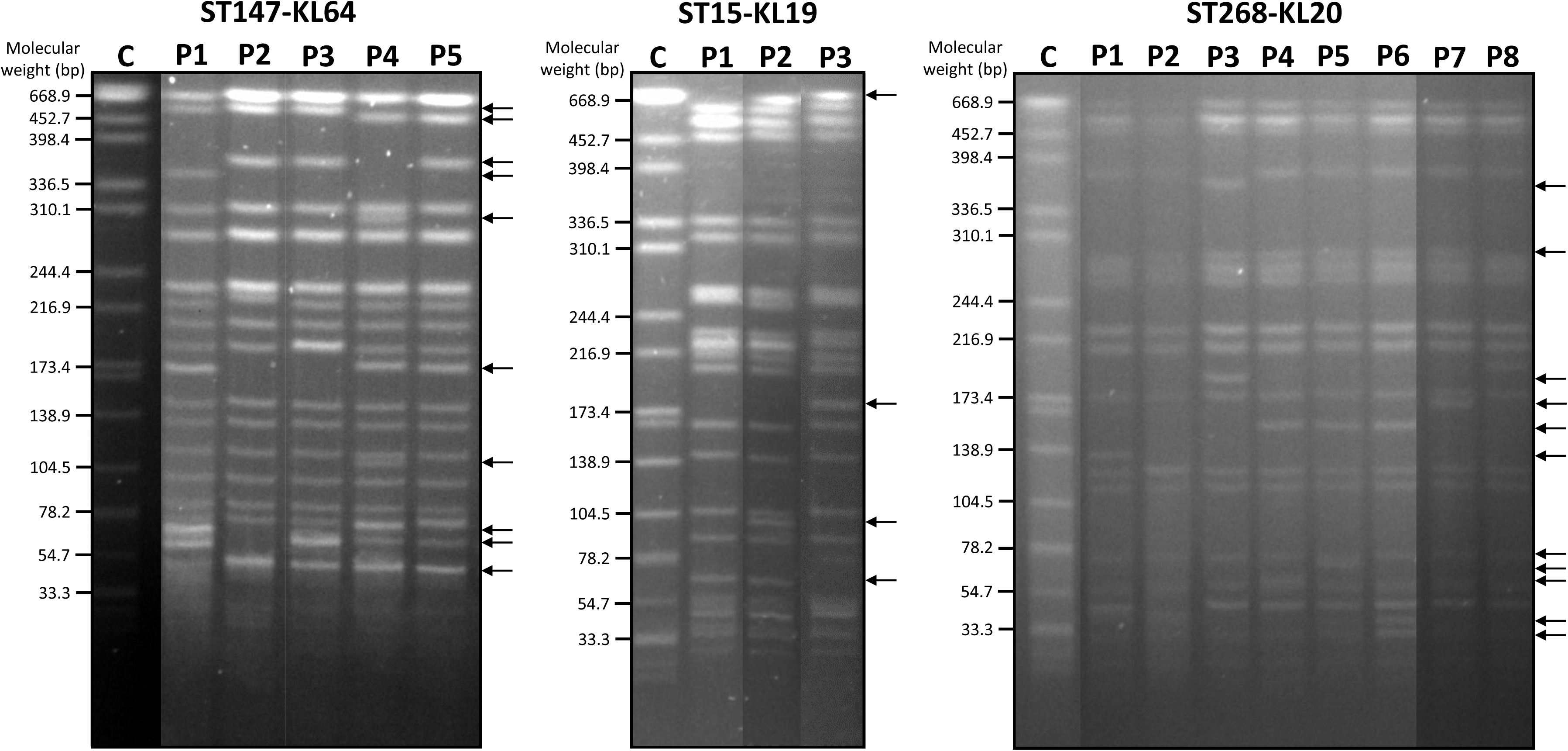
Pulsed-field gel electrophoresis (PFGE) profiles of representative isolates of the three predominant clones digested with XbaI. C represents the molecular weight control (*Salmonella enterica* serotype Braenderup H9812). The other lanes show the different *Xba*I-PFGE profiles (P) observed for the three predominant clones (ST147-KL64, ST15-KL19, and ST268-KL20). Arrows in the right represent all the differences detected in band profile within each clone group.

**Supplementary Figure 3.**
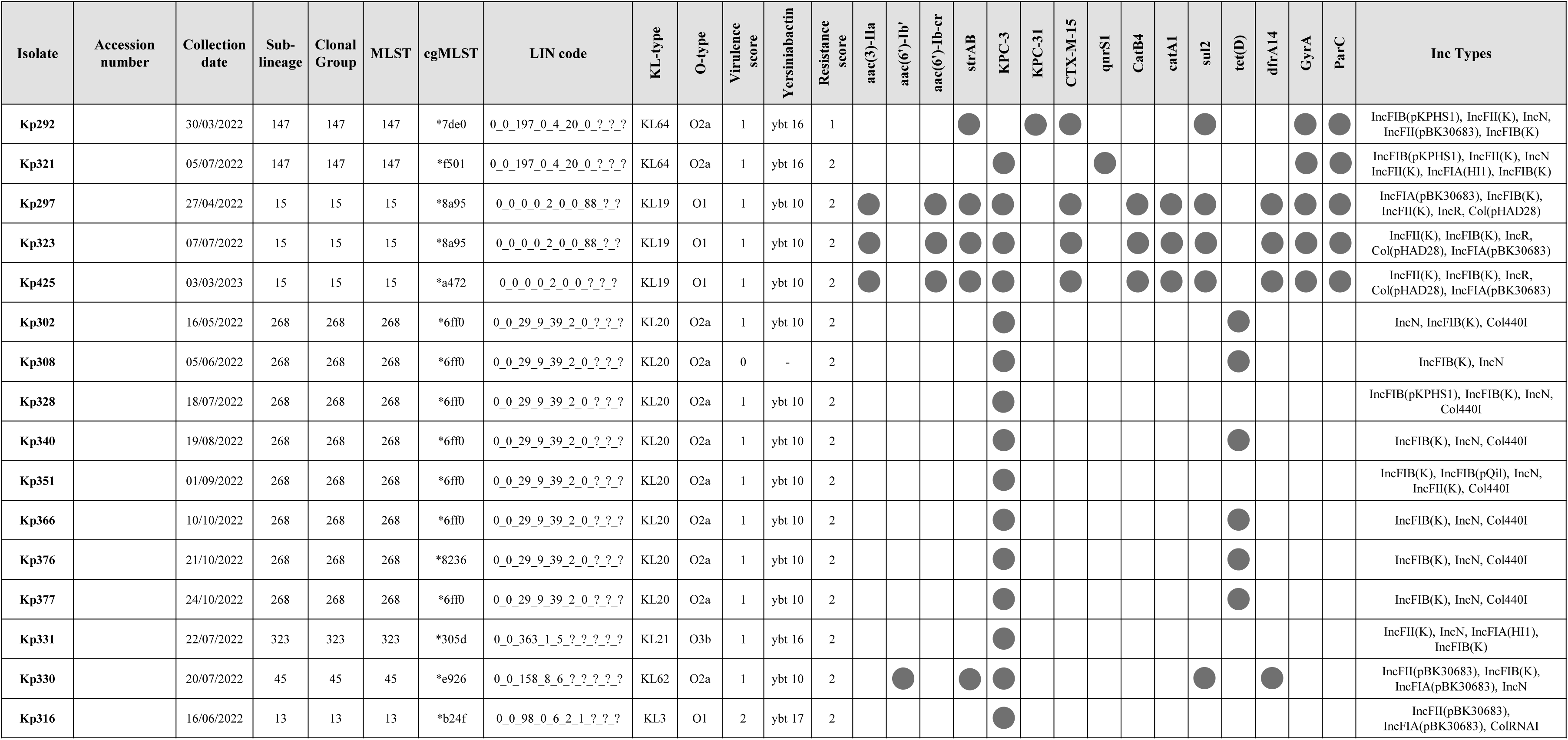
Antimicrobial resistance, virulence and replicon gene content of representative isolates sequenced by WGS obtained using Pathogenwatch.

## Supplementary Tables

**Supplementary Table S1.**
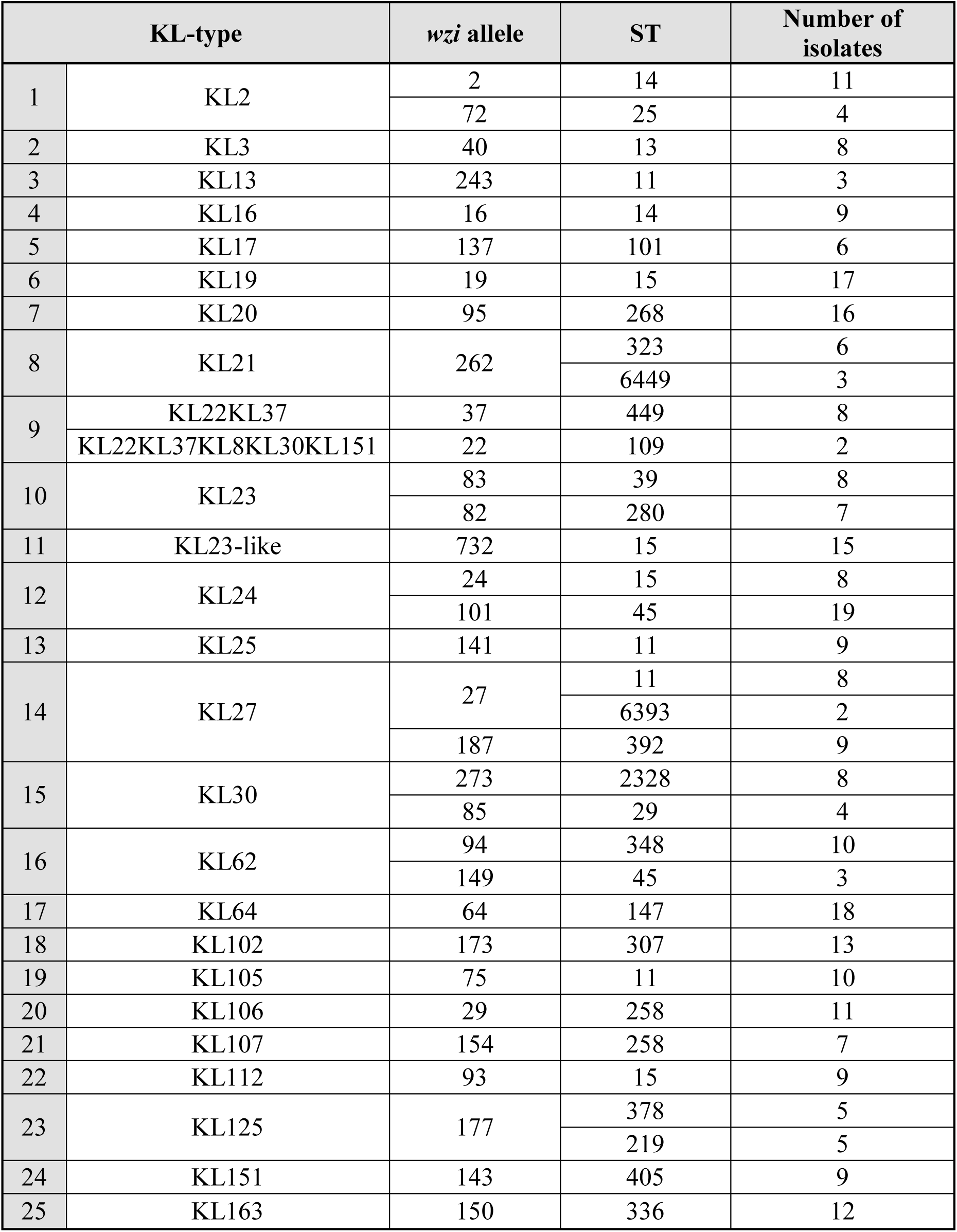
KL-types included in the training dataset for RF Model 2.

|  | KL-type | *wzi* allele | ST | Number of isolates |
| --- | --- | --- | --- | --- |
| 1 | KL2 | 2 | 14 | 11 |
|  |  | 72 | 25 | 4 |
| 2 | KL3 | 40 | 13 | 8 |
| 3 | KL13 | 243 | 11 | 3 |
| 4 | KL16 | 16 | 14 | 9 |
| 5 | KL17 | 137 | 101 | 6 |
| 6 | KL19 | 19 | 15 | 17 |
| 7 | KL20 | 95 | 268 | 16 |
| 8 | KL21 | 262 | 323 | 6 |
|  |  |  | 6449 | 3 |
| 9 | KL22KL37 | 37 | 449 | 8 |
|  | KL22KL37KL8KL30KL151 | 22 | 109 | 2 |
| 10 | KL23 | 83 | 39 | 8 |
|  |  | 82 | 280 | 7 |
| 11 | KL23-like | 732 | 15 | 15 |
| 12 | KL24 | 24 | 15 | 8 |
|  |  | 101 | 45 | 19 |
| 13 | KL25 | 141 | 11 | 9 |
| 14 | KL27 | 27 | 11 | 8 |
|  |  |  | 6393 | 2 |
|  |  | 187 | 392 | 9 |
| 15 | KL30 | 273 | 2328 | 8 |
|  |  | 85 | 29 | 4 |
| 16 | KL62 | 94 | 348 | 10 |
|  |  | 149 | 45 | 3 |
| 17 | KL64 | 64 | 147 | 18 |
| 18 | KL102 | 173 | 307 | 13 |
| 19 | KL105 | 75 | 11 | 10 |
| 20 | KL106 | 29 | 258 | 11 |
| 21 | KL107 | 154 | 258 | 7 |
| 22 | KL112 | 93 | 15 | 9 |
| 23 | KL125 | 177 | 378 | 5 |
|  |  |  | 219 | 5 |
| 24 | KL151 | 143 | 405 | 9 |
| 25 | KL163 | 150 | 336 | 12 |

**Supplementary Table S2.**
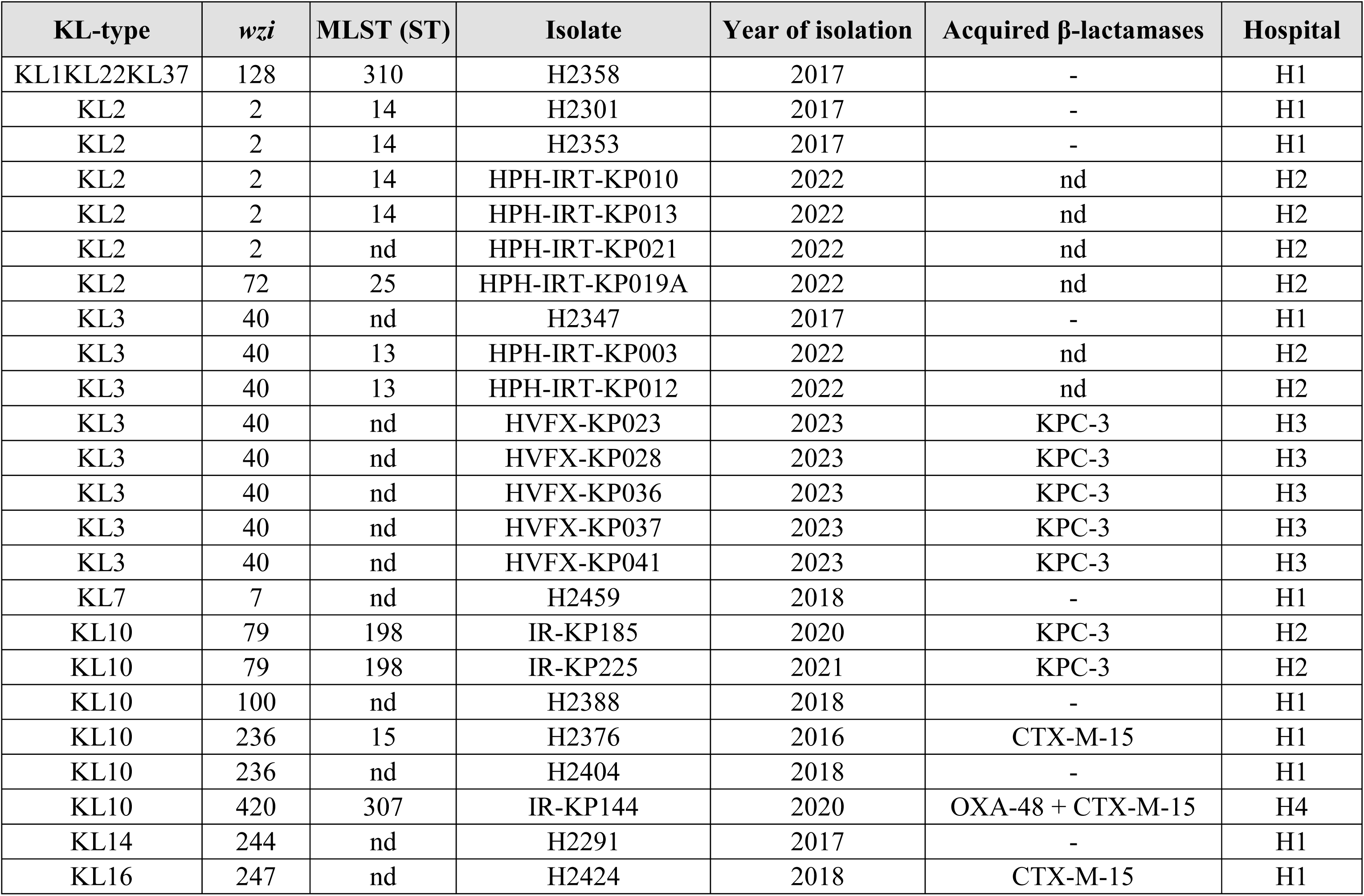
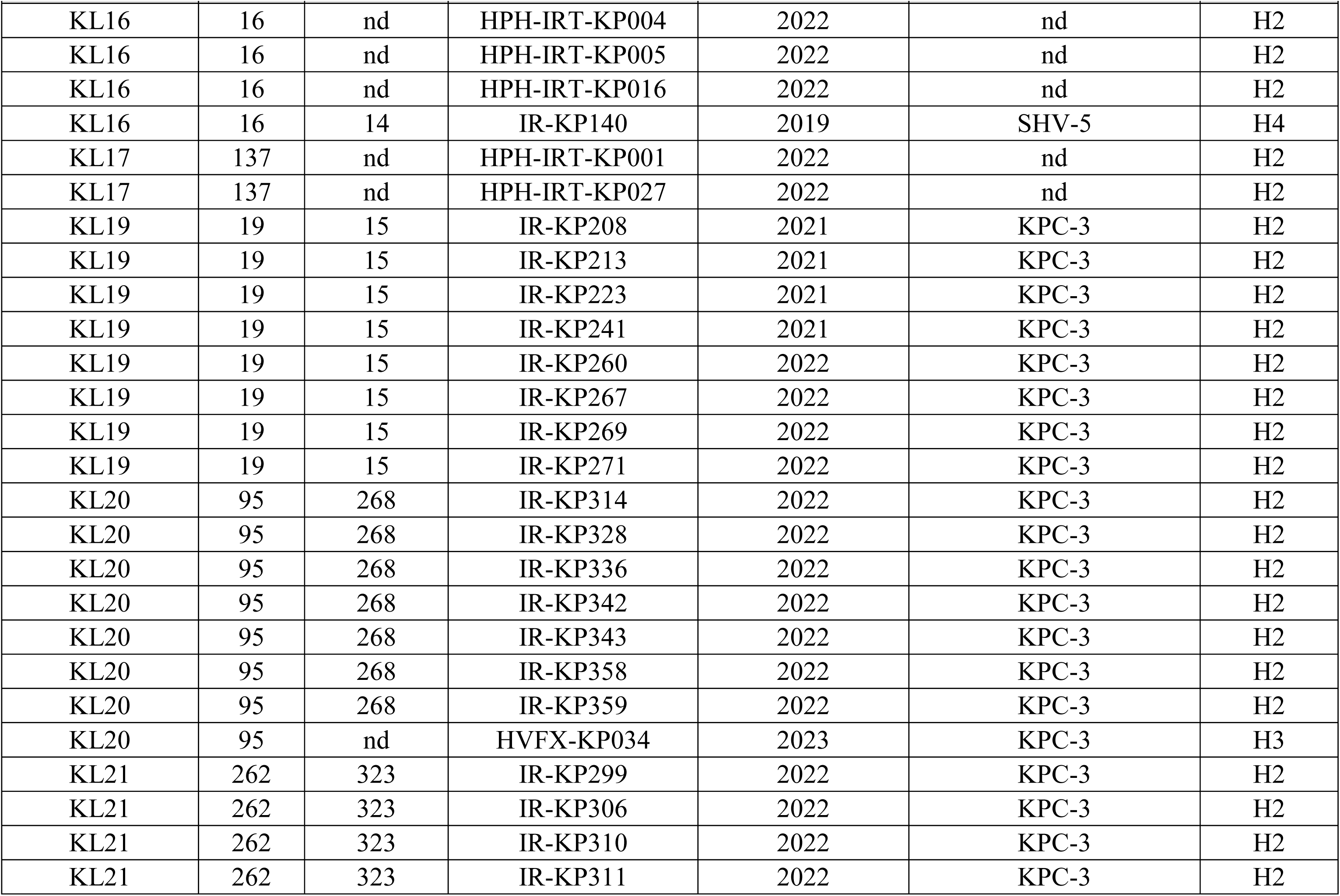
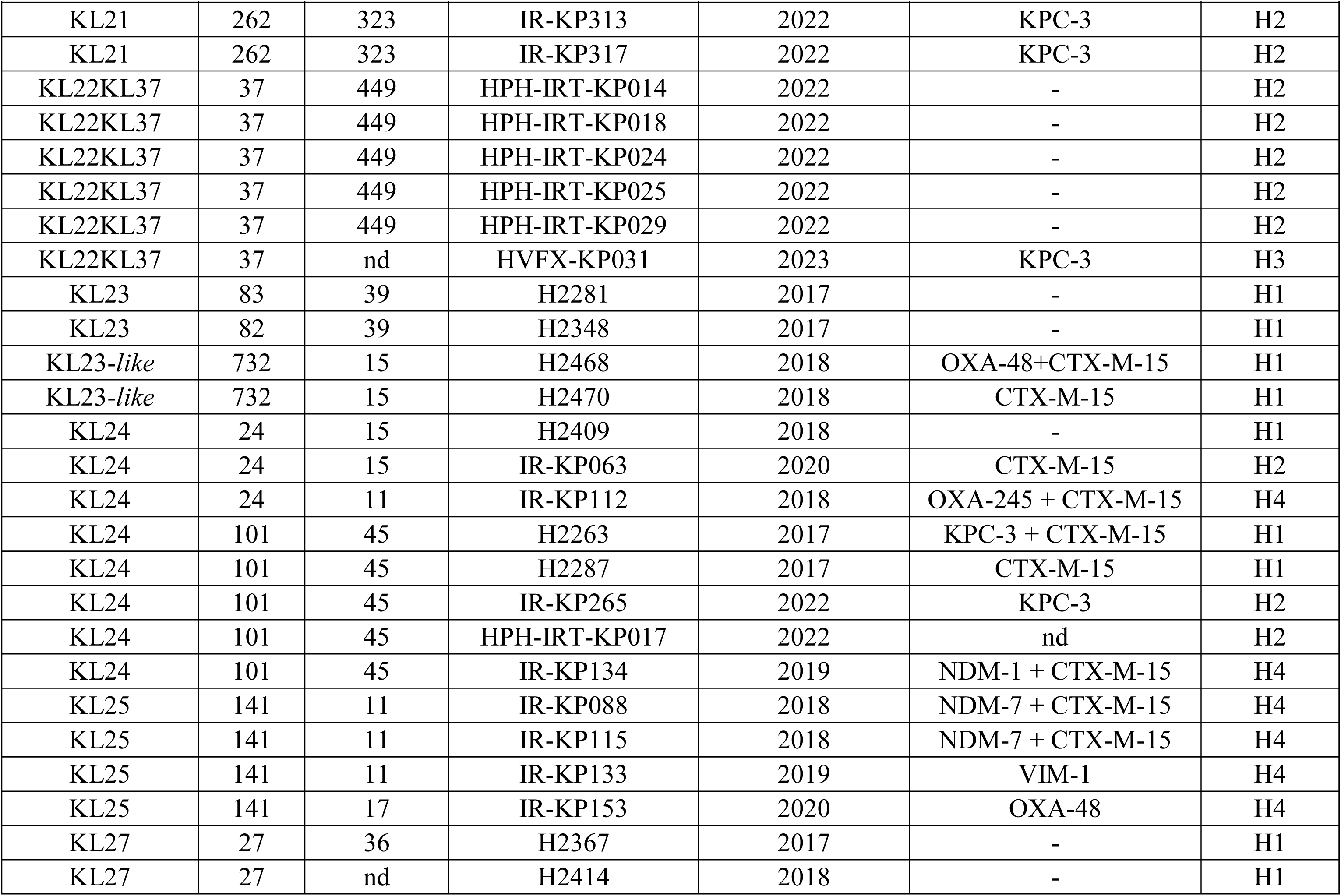
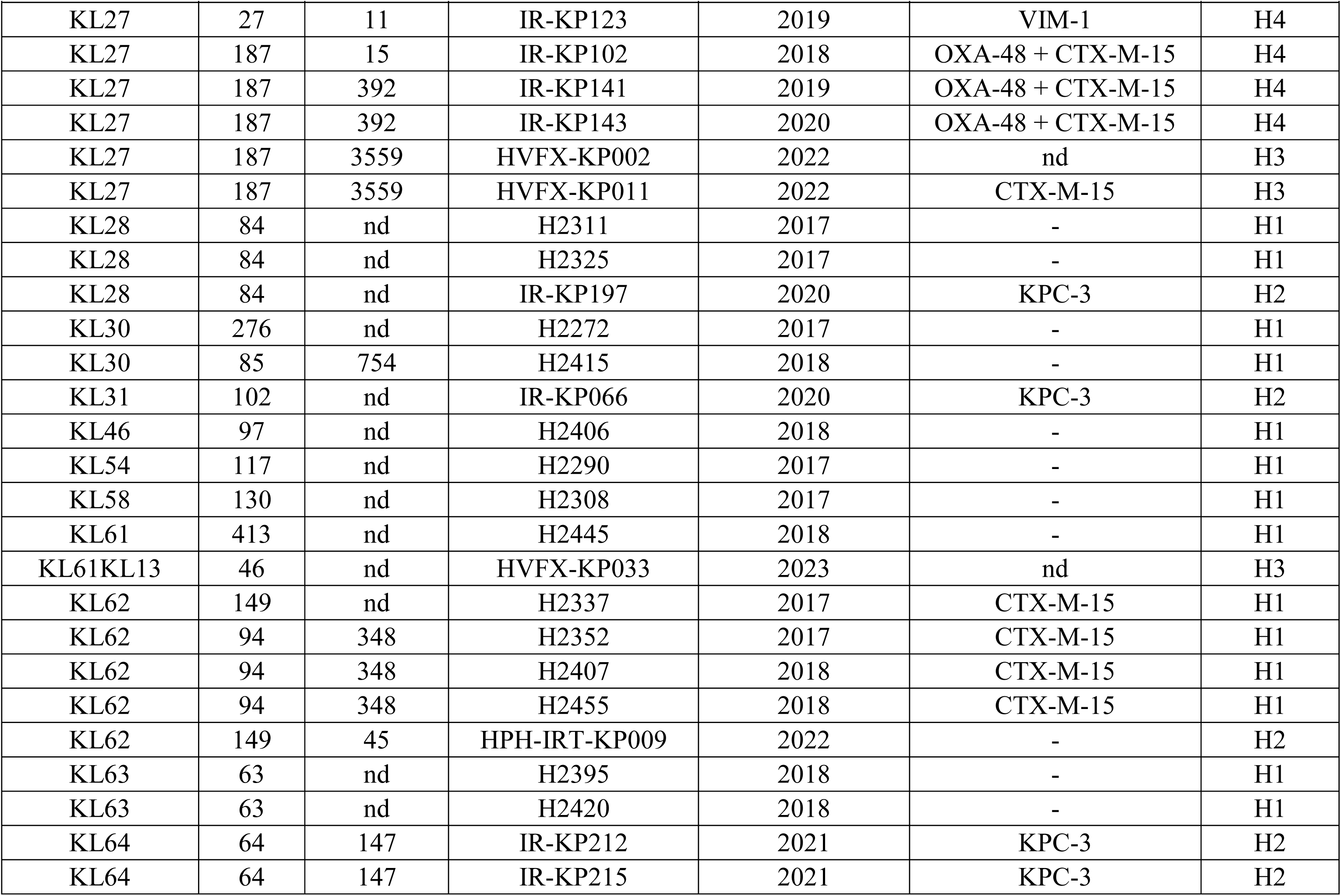
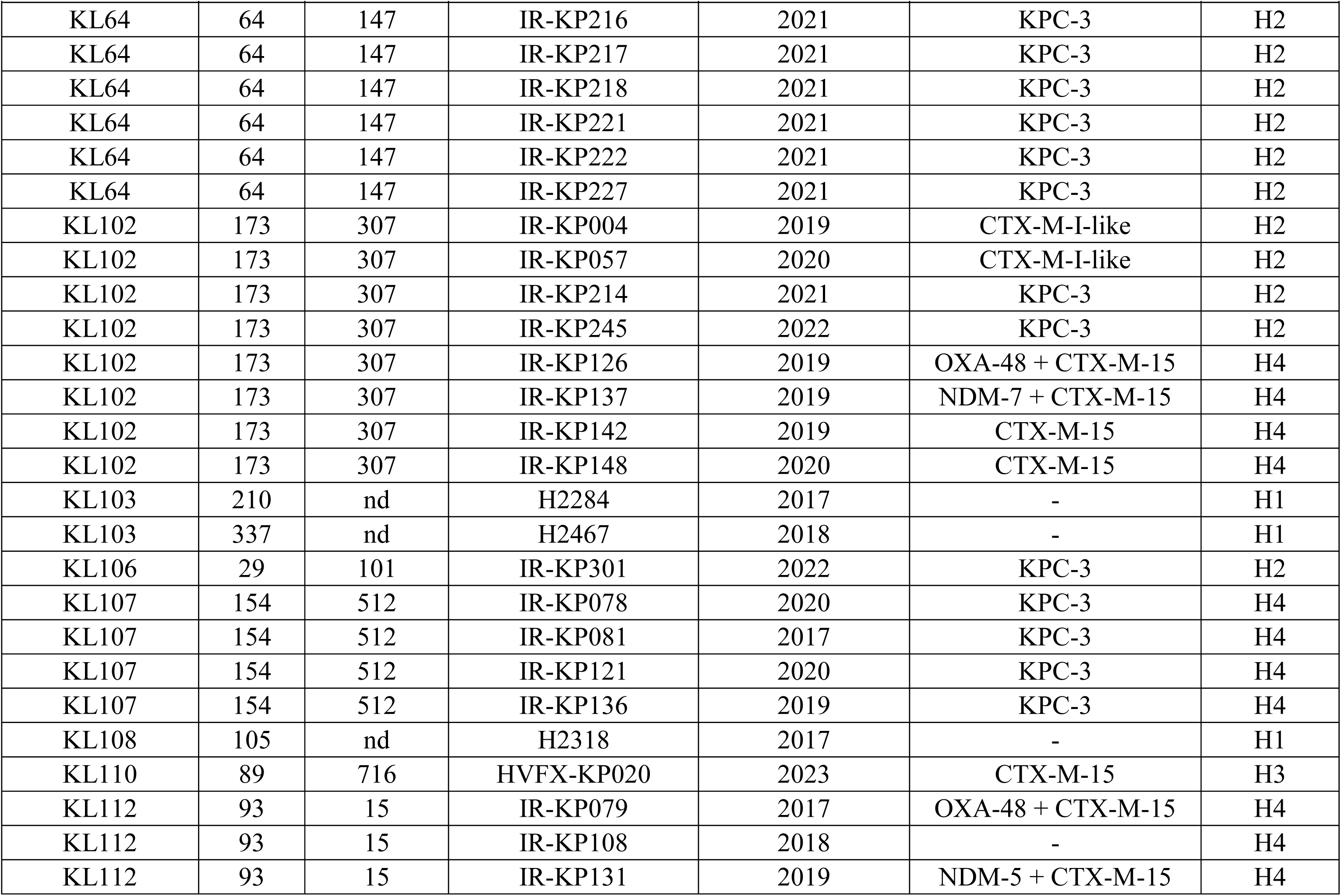
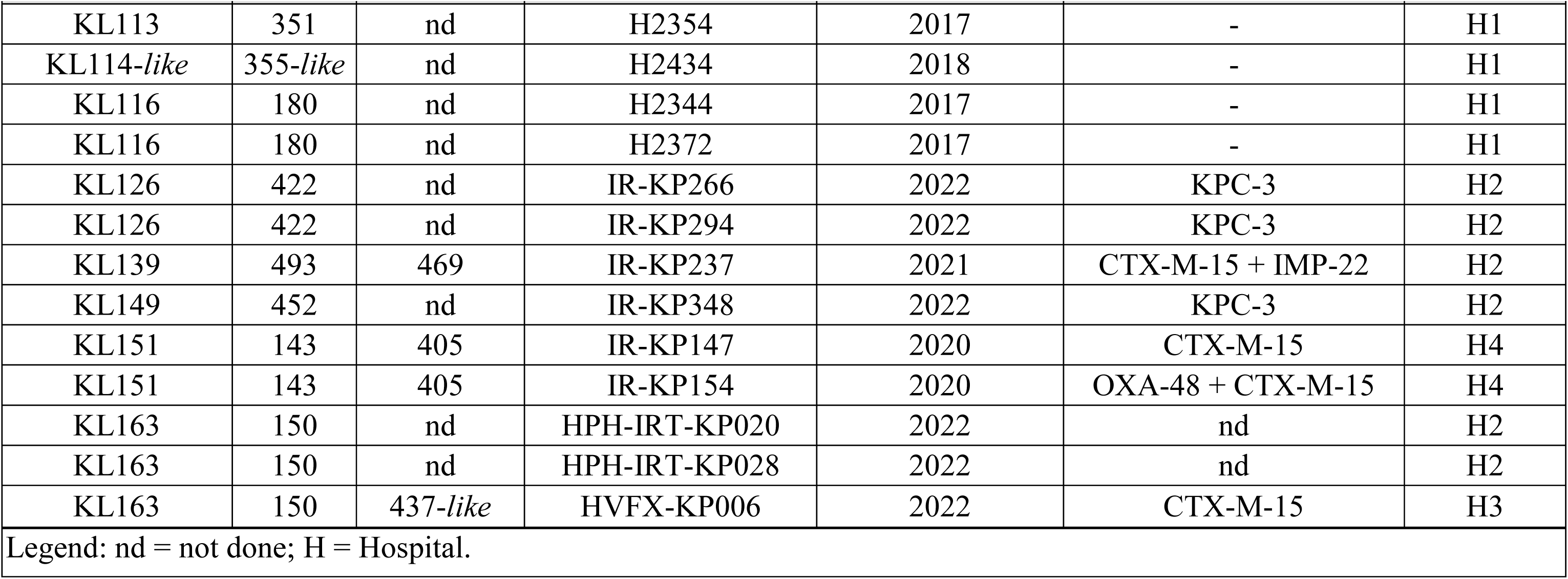
Epidemiological and genotypic data on the training set used to build the RF Model 2.

| KL-type | *wzi* | MLST (ST) | Isolate | Year of isolation | Acquired β-lactamases | Hospital |
| --- | --- | --- | --- | --- | --- | --- |
| KL1KL22KL37 | 128 | 310 | H2358 | 2017 | - | H1 |
| KL2 | 2 | 14 | H2301 | 2017 | - | H1 |
| KL2 | 2 | 14 | H2353 | 2017 | - | H1 |
| KL2 | 2 | 14 | HPH-IRT-KP010 | 2022 | nd | H2 |
| KL2 | 2 | 14 | HPH-IRT-KP013 | 2022 | nd | H2 |
| KL2 | 2 | nd | HPH-IRT-KP021 | 2022 | nd | H2 |
| KL2 | 72 | 25 | HPH-IRT-KP019A | 2022 | nd | H2 |
| KL3 | 40 | nd | H2347 | 2017 | - | H1 |
| KL3 | 40 | 13 | HPH-IRT-KP003 | 2022 | nd | H2 |
| KL3 | 40 | 13 | HPH-IRT-KP012 | 2022 | nd | H2 |
| KL3 | 40 | nd | HVFX-KP023 | 2023 | KPC-3 | H3 |
| KL3 | 40 | nd | HVFX-KP028 | 2023 | KPC-3 | H3 |
| KL3 | 40 | nd | HVFX-KP036 | 2023 | KPC-3 | H3 |
| KL3 | 40 | nd | HVFX-KP037 | 2023 | KPC-3 | H3 |
| KL3 | 40 | nd | HVFX-KP041 | 2023 | KPC-3 | H3 |
| KL7 | 7 | nd | H2459 | 2018 | - | H1 |
| KL10 | 79 | 198 | IR-KP185 | 2020 | KPC-3 | H2 |
| KL10 | 79 | 198 | IR-KP225 | 2021 | KPC-3 | H2 |
| KL10 | 100 | nd | H2388 | 2018 | - | H1 |
| KL10 | 236 | 15 | H2376 | 2016 | CTX-M-15 | H1 |
| KL10 | 236 | nd | H2404 | 2018 | - | H1 |
| KL10 | 420 | 307 | IR-KP144 | 2020 | OXA-48 + CTX-M-15 | H4 |
| KL14 | 244 | nd | H2291 | 2017 | - | H1 |
| KL16 | 247 | nd | H2424 | 2018 | CTX-M-15 | H1 |
| KL16 | 16 | nd | HPH-IRT-KP004 | 2022 | nd | H2 |
| KL16 | 16 | nd | HPH-IRT-KP005 | 2022 | nd | H2 |
| KL16 | 16 | nd | HPH-IRT-KP016 | 2022 | nd | H2 |
| KL16 | 16 | 14 | IR-KP140 | 2019 | SHV-5 | H4 |
| KL17 | 137 | nd | HPH-IRT-KP001 | 2022 | nd | H2 |
| KL17 | 137 | nd | HPH-IRT-KP027 | 2022 | nd | H2 |
| KL19 | 19 | 15 | IR-KP208 | 2021 | KPC-3 | H2 |
| KL19 | 19 | 15 | IR-KP213 | 2021 | KPC-3 | H2 |
| KL19 | 19 | 15 | IR-KP223 | 2021 | KPC-3 | H2 |
| KL19 | 19 | 15 | IR-KP241 | 2021 | KPC-3 | H2 |
| KL19 | 19 | 15 | IR-KP260 | 2022 | KPC-3 | H2 |
| KL19 | 19 | 15 | IR-KP267 | 2022 | KPC-3 | H2 |
| KL19 | 19 | 15 | IR-KP269 | 2022 | KPC-3 | H2 |
| KL19 | 19 | 15 | IR-KP271 | 2022 | KPC-3 | H2 |
| KL20 | 95 | 268 | IR-KP314 | 2022 | KPC-3 | H2 |
| KL20 | 95 | 268 | IR-KP328 | 2022 | KPC-3 | H2 |
| KL20 | 95 | 268 | IR-KP336 | 2022 | KPC-3 | H2 |
| KL20 | 95 | 268 | IR-KP342 | 2022 | KPC-3 | H2 |
| KL20 | 95 | 268 | IR-KP343 | 2022 | KPC-3 | H2 |
| KL20 | 95 | 268 | IR-KP358 | 2022 | KPC-3 | H2 |
| KL20 | 95 | 268 | IR-KP359 | 2022 | KPC-3 | H2 |
| KL20 | 95 | nd | HVFX-KP034 | 2023 | KPC-3 | H3 |
| KL21 | 262 | 323 | IR-KP299 | 2022 | KPC-3 | H2 |
| KL21 | 262 | 323 | IR-KP306 | 2022 | KPC-3 | H2 |
| KL21 | 262 | 323 | IR-KP310 | 2022 | KPC-3 | H2 |
| KL21 | 262 | 323 | IR-KP311 | 2022 | KPC-3 | H2 |

| KL-type | wzi | MLST (ST) | Isolate | Year of isolation | Acquired β-lactamases | Hospital |
| --- | --- | --- | --- | --- | --- | --- |
| KL21 | 262 | 323 | IR-KP313 | 2022 | KPC-3 | H2 |
| KL21 | 262 | 323 | IR-KP317 | 2022 | KPC-3 | H2 |
| KL22KL37 | 37 | 449 | HPH-IRT-KP014 | 2022 | - | H2 |
| KL22KL37 | 37 | 449 | HPH-IRT-KP018 | 2022 | - | H2 |
| KL22KL37 | 37 | 449 | HPH-IRT-KP024 | 2022 | - | H2 |
| KL22KL37 | 37 | 449 | HPH-IRT-KP025 | 2022 | - | H2 |
| KL22KL37 | 37 | 449 | HPH-IRT-KP029 | 2022 | - | H2 |
| KL22KL37 | 37 | nd | HVFX-KP031 | 2023 | KPC-3 | H3 |
| KL23 | 83 | 39 | H2281 | 2017 | - | H1 |
| KL23 | 82 | 39 | H2348 | 2017 | - | H1 |
| KL23-like | 732 | 15 | H2468 | 2018 | OXA-48+CTX-M-15 | H1 |
| KL23-like | 732 | 15 | H2470 | 2018 | CTX-M-15 | H1 |
| KL24 | 24 | 15 | H2409 | 2018 | - | H1 |
| KL24 | 24 | 15 | IR-KP063 | 2020 | CTX-M-15 | H2 |
| KL24 | 24 | 11 | IR-KP112 | 2018 | OXA-245 + CTX-M-15 | H4 |
| KL24 | 101 | 45 | H2263 | 2017 | KPC-3 + CTX-M-15 | H1 |
| KL24 | 101 | 45 | H2287 | 2017 | CTX-M-15 | H1 |
| KL24 | 101 | 45 | IR-KP265 | 2022 | KPC-3 | H2 |
| KL24 | 101 | 45 | HPH-IRT-KP017 | 2022 | nd | H2 |
| KL24 | 101 | 45 | IR-KP134 | 2019 | NDM-1 + CTX-M-15 | H4 |
| KL25 | 141 | 11 | IR-KP088 | 2018 | NDM-7 + CTX-M-15 | H4 |
| KL25 | 141 | 11 | IR-KP115 | 2018 | NDM-7 + CTX-M-15 | H4 |
| KL25 | 141 | 11 | IR-KP133 | 2019 | VIM-1 | H4 |
| KL25 | 141 | 17 | IR-KP153 | 2020 | OXA-48 | H4 |
| KL27 | 27 | 36 | H2367 | 2017 | - | H1 |
| KL27 | 27 | nd | H2414 | 2018 | - | H1 |
| KL27 | 27 | 11 | IR-KP123 | 2019 | VIM-1 | H4 |
| KL27 | 187 | 15 | IR-KP102 | 2018 | OXA-48 + CTX-M-15 | H4 |
| KL27 | 187 | 392 | IR-KP141 | 2019 | OXA-48 + CTX-M-15 | H4 |
| KL27 | 187 | 392 | IR-KP143 | 2020 | OXA-48 + CTX-M-15 | H4 |
| KL27 | 187 | 3559 | HVFX-KP002 | 2022 | nd | H3 |
| KL27 | 187 | 3559 | HVFX-KP011 | 2022 | CTX-M-15 | H3 |
| KL28 | 84 | nd | H2311 | 2017 | - | H1 |
| KL28 | 84 | nd | H2325 | 2017 | - | H1 |
| KL28 | 84 | nd | IR-KP197 | 2020 | KPC-3 | H2 |
| KL30 | 276 | nd | H2272 | 2017 | - | H1 |
| KL30 | 85 | 754 | H2415 | 2018 | - | H1 |
| KL31 | 102 | nd | IR-KP066 | 2020 | KPC-3 | H2 |
| KL46 | 97 | nd | H2406 | 2018 | - | H1 |
| KL54 | 117 | nd | H2290 | 2017 | - | H1 |
| KL58 | 130 | nd | H2308 | 2017 | - | H1 |
| KL61 | 413 | nd | H2445 | 2018 | - | H1 |
| KL61KL13 | 46 | nd | HVFX-KP033 | 2023 | nd | H3 |
| KL62 | 149 | nd | H2337 | 2017 | CTX-M-15 | H1 |
| KL62 | 94 | 348 | H2352 | 2017 | CTX-M-15 | H1 |
| KL62 | 94 | 348 | H2407 | 2018 | CTX-M-15 | H1 |
| KL62 | 94 | 348 | H2455 | 2018 | CTX-M-15 | H1 |
| KL62 | 149 | 45 | HPH-IRT-KP009 | 2022 | - | H2 |
| KL63 | 63 | nd | H2395 | 2018 | - | H1 |
| KL63 | 63 | nd | H2420 | 2018 | - | H1 |
| KL64 | 64 | 147 | IR-KP212 | 2021 | KPC-3 | H2 |
| KL64 | 64 | 147 | IR-KP215 | 2021 | KPC-3 | H2 |
| KL64 | 64 | 147 | IR-KP216 | 2021 | KPC-3 | H2 |
| KL64 | 64 | 147 | IR-KP217 | 2021 | KPC-3 | H2 |
| KL64 | 64 | 147 | IR-KP218 | 2021 | KPC-3 | H2 |
| KL64 | 64 | 147 | IR-KP221 | 2021 | KPC-3 | H2 |
| KL64 | 64 | 147 | IR-KP222 | 2021 | KPC-3 | H2 |
| KL64 | 64 | 147 | IR-KP227 | 2021 | KPC-3 | H2 |
| KL102 | 173 | 307 | IR-KP004 | 2019 | CTX-M-I-like | H2 |
| KL102 | 173 | 307 | IR-KP057 | 2020 | CTX-M-I-like | H2 |
| KL102 | 173 | 307 | IR-KP214 | 2021 | KPC-3 | H2 |
| KL102 | 173 | 307 | IR-KP245 | 2022 | KPC-3 | H2 |
| KL102 | 173 | 307 | IR-KP126 | 2019 | OXA-48 + CTX-M-15 | H4 |
| KL102 | 173 | 307 | IR-KP137 | 2019 | NDM-7 + CTX-M-15 | H4 |
| KL102 | 173 | 307 | IR-KP142 | 2019 | CTX-M-15 | H4 |
| KL102 | 173 | 307 | IR-KP148 | 2020 | CTX-M-15 | H4 |
| KL103 | 210 | nd | H2284 | 2017 | - | H1 |
| KL103 | 337 | nd | H2467 | 2018 | - | H1 |
| KL106 | 29 | 101 | IR-KP301 | 2022 | KPC-3 | H2 |
| KL107 | 154 | 512 | IR-KP078 | 2020 | KPC-3 | H4 |
| KL107 | 154 | 512 | IR-KP081 | 2017 | KPC-3 | H4 |
| KL107 | 154 | 512 | IR-KP121 | 2020 | KPC-3 | H4 |
| KL107 | 154 | 512 | IR-KP136 | 2019 | KPC-3 | H4 |
| KL108 | 105 | nd | H2318 | 2017 | - | H1 |
| KL110 | 89 | 716 | HVFX-KP020 | 2023 | CTX-M-15 | H3 |
| KL112 | 93 | 15 | IR-KP079 | 2017 | OXA-48 + CTX-M-15 | H4 |
| KL112 | 93 | 15 | IR-KP108 | 2018 | - | H4 |
| KL112 | 93 | 15 | IR-KP131 | 2019 | NDM-5 + CTX-M-15 | H4 |
| KL113 | 351 | nd | H2354 | 2017 | - | H1 |
| KL114-like | 355-like | nd | H2434 | 2018 | - | H1 |
| KL116 | 180 | nd | H2344 | 2017 | - | H1 |
| KL116 | 180 | nd | H2372 | 2017 | - | H1 |
| KL126 | 422 | nd | IR-KP266 | 2022 | KPC-3 | H2 |
| KL126 | 422 | nd | IR-KP294 | 2022 | KPC-3 | H2 |
| KL139 | 493 | 469 | IR-KP237 | 2021 | CTX-M-15 + IMP-22 | H2 |
| KL149 | 452 | nd | IR-KP348 | 2022 | KPC-3 | H2 |
| KL151 | 143 | 405 | IR-KP147 | 2020 | CTX-M-15 | H4 |
| KL151 | 143 | 405 | IR-KP154 | 2020 | OXA-48 + CTX-M-15 | H4 |
| KL163 | 150 | nd | HPH-IRT-KP020 | 2022 | nd | H2 |
| KL163 | 150 | nd | HPH-IRT-KP028 | 2022 | nd | H2 |
| KL163 | 150 | 437-like | HVFX-KP006 | 2022 | CTX-M-15 | H3 |
| Legend: nd = not done; H = Hospital. |  |  |  |  |  |  |

**Supplementary Table S3.**
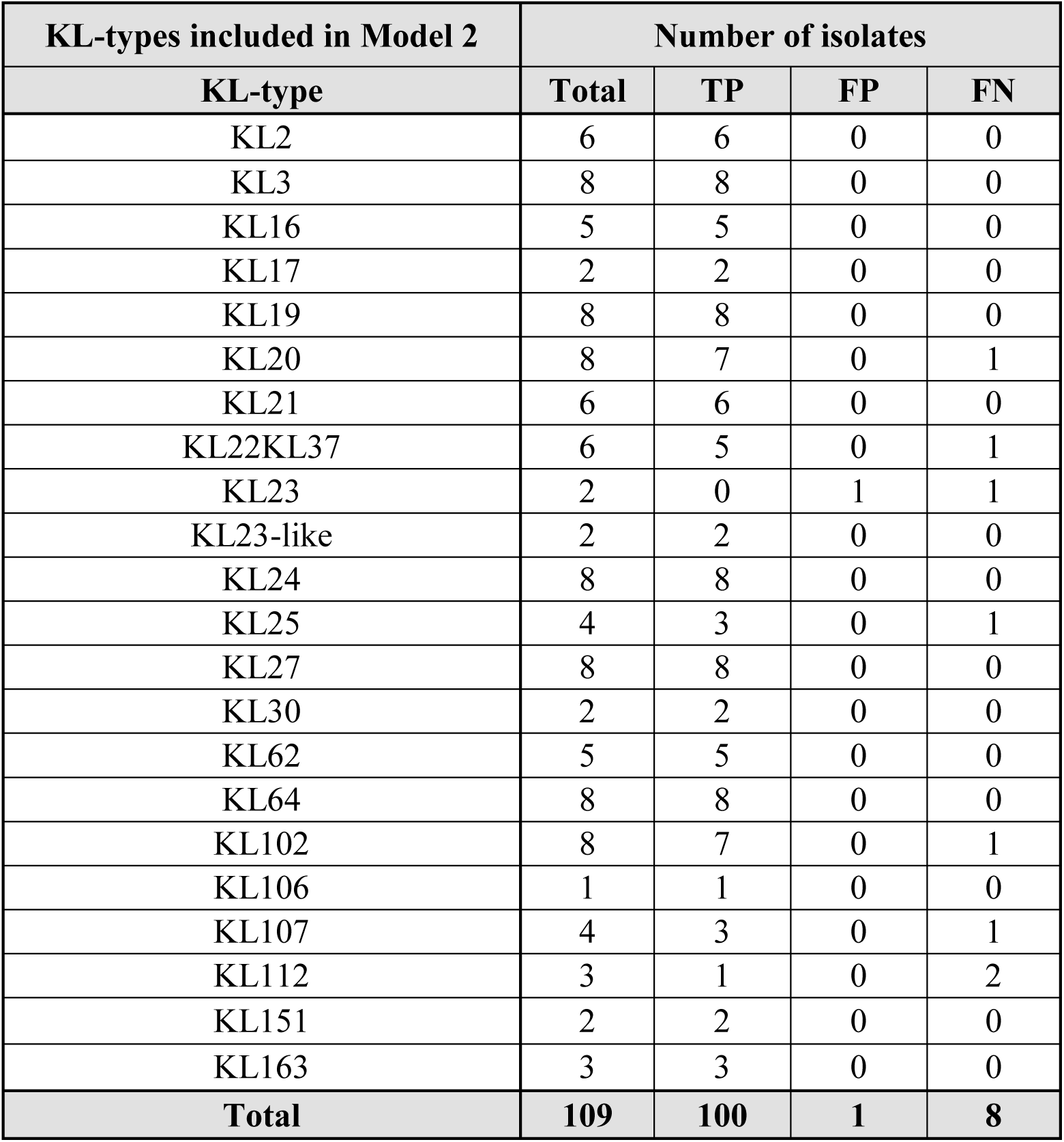
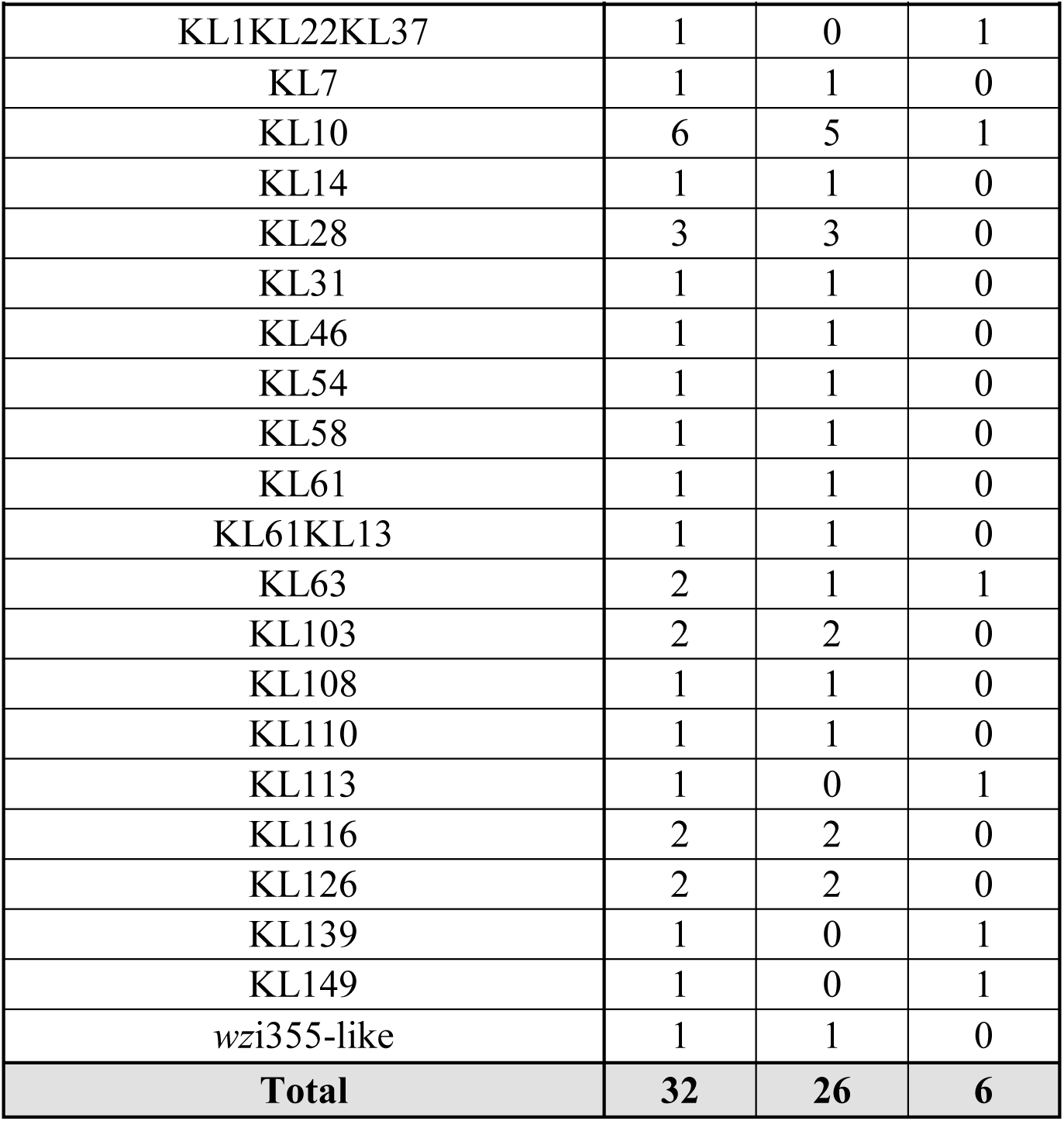
Validation set and performance standards obtained with the RF Model 2.

| **KL-types included in Model 2** | **Number of isolates** |  |  |  |
| --- | --- | --- | --- | --- |
| **KL-type** | **Total** | **TP** | **FP** | **FN** |
| KL2 | 6 | 6 | 0 | 0 |
| KL3 | 8 | 8 | 0 | 0 |
| KL16 | 5 | 5 | 0 | 0 |
| KL17 | 2 | 2 | 0 | 0 |
| KL19 | 8 | 8 | 0 | 0 |
| KL20 | 8 | 7 | 0 | 1 |
| KL21 | 6 | 6 | 0 | 0 |
| KL22KL37 | 6 | 5 | 0 | 1 |
| KL23 | 2 | 0 | 1 | 1 |
| KL23-like | 2 | 2 | 0 | 0 |
| KL24 | 8 | 8 | 0 | 0 |
| KL25 | 4 | 3 | 0 | 1 |
| KL27 | 8 | 8 | 0 | 0 |
| KL30 | 2 | 2 | 0 | 0 |
| KL62 | 5 | 5 | 0 | 0 |
| KL64 | 8 | 8 | 0 | 0 |
| KL102 | 8 | 7 | 0 | 1 |
| KL106 | 1 | 1 | 0 | 0 |
| KL107 | 4 | 3 | 0 | 1 |
| KL112 | 3 | 1 | 0 | 2 |
| KL151 | 2 | 2 | 0 | 0 |
| KL163 | 3 | 3 | 0 | 0 |
| **Total** | **109** | **100** | **1** | **8** |

| **KL-types not included in Model 2** | **Number of isolates** |  |  |
| --- | --- | --- | --- |
| **KL-type** | **Total** | **TN** | **FP** |
| KL1KL22KL37 | 1 | 0 | 1 |
| KL7 | 1 | 1 | 0 |
| KL10 | 6 | 5 | 1 |
| KL14 | 1 | 1 | 0 |
| KL28 | 3 | 3 | 0 |
| KL31 | 1 | 1 | 0 |
| KL46 | 1 | 1 | 0 |
| KL54 | 1 | 1 | 0 |
| KL58 | 1 | 1 | 0 |
| KL61 | 1 | 1 | 0 |
| KL61KL13 | 1 | 1 | 0 |
| KL63 | 2 | 1 | 1 |
| KL103 | 2 | 2 | 0 |
| KL108 | 1 | 1 | 0 |
| KL110 | 1 | 1 | 0 |
| KL113 | 1 | 0 | 1 |
| KL116 | 2 | 2 | 0 |
| KL126 | 2 | 2 | 0 |
| KL139 | 1 | 0 | 1 |
| KL149 | 1 | 0 | 1 |
| wzi355-like | 1 | 1 | 0 |
| **Total** | **32** | **26** | **6** |

**Supplementary Table S4.** Pairwise single nucleotide polymorphism distances score matrix of ST268-KL20 core genome-based neighbour-joining phylogenetic tree obtained in Pathogenwatch.

| Isolate |  | Isolates from this study |  |  |  |  |  |  |  | Isolates from Pathogenwatch database |  |  |  |  |  |  |  |  |  |  |  |  |  |  |  |
| --- | --- | --- | --- | --- | --- | --- | --- | --- | --- | --- | --- | --- | --- | --- | --- | --- | --- | --- | --- | --- | --- | --- | --- | --- | --- |
|  |  | Kp302 | Kp308 | Kp377 | Kp340 | Kp351 | Kp328 | Kp376 | Kp366 | SAMN34998335 | SRR12902472 | SAMN33905359 | SRR8844730 | SAMN35130898 | SAMN26976245 | SRR11819369 | SRR11819368 | SRR14450288 | SRR14449659 | SRR12509575 | SRR8199632 | SRR8198917 | SRR17431323 | SAMN34078859 | SRR16847985 |
| Isolates from this study | Kp308 | 4 |  |  |  |  |  |  |  |  |  |  |  |  |  |  |  |  |  |  |  |  |  |  |  |
|  | Kp377 | 2 | 4 |  |  |  |  |  |  |  |  |  |  |  |  |  |  |  |  |  |  |  |  |  |  |
|  | Kp340 | 2 | 4 | 2 |  |  |  |  |  |  |  |  |  |  |  |  |  |  |  |  |  |  |  |  |  |
|  | Kp351 | 3 | 5 | 3 | 3 |  |  |  |  |  |  |  |  |  |  |  |  |  |  |  |  |  |  |  |  |
|  | Kp328 | 1 | 3 | 1 | 1 | 2 |  |  |  |  |  |  |  |  |  |  |  |  |  |  |  |  |  |  |  |
|  | Kp376 | 3 | 5 | 3 | 3 | 4 | 2 |  |  |  |  |  |  |  |  |  |  |  |  |  |  |  |  |  |  |
|  | Kp366 | 2 | 4 | 2 | 2 | 3 | 1 | 1 |  |  |  |  |  |  |  |  |  |  |  |  |  |  |  |  |  |
| Isolates from Pathogenwatch database | SAMN34998335 | 22 | 24 | 22 | 22 | 23 | 21 | 23 | 22 |  |  |  |  |  |  |  |  |  |  |  |  |  |  |  |  |
|  | SRR12902472 | 12 | 14 | 12 | 12 | 13 | 11 | 13 | 12 | 13 |  |  |  |  |  |  |  |  |  |  |  |  |  |  |  |
|  | SAMN33905359 | 22 | 24 | 22 | 22 | 23 | 21 | 23 | 22 | 23 | 13 |  |  |  |  |  |  |  |  |  |  |  |  |  |  |
|  | SRR8844730 | 20 | 22 | 20 | 20 | 21 | 19 | 21 | 20 | 21 | 11 | 21 |  |  |  |  |  |  |  |  |  |  |  |  |  |
|  | SAMN35130898 | 26 | 28 | 26 | 26 | 27 | 25 | 27 | 26 | 27 | 18 | 27 | 25 |  |  |  |  |  |  |  |  |  |  |  |  |
|  | SAMN26976245 | 21 | 23 | 21 | 21 | 22 | 20 | 22 | 21 | 22 | 12 | 22 | 20 | 15 |  |  |  |  |  |  |  |  |  |  |  |
|  | SRR11819369 | 65 | 67 | 65 | 65 | 66 | 64 | 66 | 65 | 66 | 57 | 66 | 64 | 70 | 65 |  |  |  |  |  |  |  |  |  |  |
|  | SRR11819368 | 64 | 66 | 64 | 64 | 65 | 63 | 65 | 64 | 65 | 56 | 65 | 63 | 69 | 64 | 1 |  |  |  |  |  |  |  |  |  |
|  |  | Kp302 | Kp308 | Kp377 | Kp340 | Kp351 | Kp328 | Kp376 | Kp366 | SAMN34998335 | SRR12902472 | SAMN33905359 | SRR8844730 | SAMN35130898 | SAMN26976245 | SRR11819369 | SRR11819368 | SRR14450288 | SRR14449659 | SRR12509575 | SRR8199632 | SRR8198917 | SRR17431323 | SAMN34078859 | SRR16847985 |
| Isolates from Pathogenwatch database | SRR14450288 | 60 | 62 | 60 | 60 | 61 | 59 | 61 | 60 | 61 | 52 | 61 | 59 | 65 | 60 | 9 | 8 |  |  |  |  |  |  |  |  |
|  | SRR14449659 | 58 | 60 | 58 | 58 | 59 | 57 | 59 | 58 | 59 | 49 | 59 | 57 | 63 | 58 | 7 | 6 | 2 |  |  |  |  |  |  |  |
|  | SRR12509575 | 63 | 65 | 63 | 63 | 64 | 62 | 64 | 63 | 64 | 55 | 64 | 62 | 68 | 63 | 14 | 13 | 9 | 7 |  |  |  |  |  |  |
|  | SRR8199632 | 71 | 74 | 71 | 71 | 72 | 70 | 72 | 71 | 73 | 63 | 72 | 70 | 76 | 72 | 31 | 30 | 26 | 24 | 29 |  |  |  |  |  |
|  | SRR8198917 | 61 | 63 | 61 | 61 | 62 | 60 | 62 | 61 | 63 | 54 | 63 | 61 | 67 | 62 | 22 | 21 | 17 | 14 | 20 | 3 |  |  |  |  |
|  | SRR17431323 | 73 | 75 | 73 | 73 | 74 | 72 | 74 | 73 | 74 | 65 | 74 | 72 | 78 | 73 | 84 | 83 | 79 | 77 | 82 | 91 | 82 |  |  |  |
|  | SAMN34078859 | 22 | 24 | 22 | 22 | 23 | 21 | 23 | 22 | 23 | 13 | 23 | 21 | 27 | 22 | 60 | 59 | 55 | 53 | 58 | 66 | 57 | 68 |  |  |
|  | SRR16847985 | 24 | 26 | 24 | 24 | 25 | 23 | 25 | 24 | 25 | 15 | 25 | 23 | 29 | 24 | 64 | 63 | 59 | 57 | 62 | 70 | 61 | 72 | 21 |  |
|  | SAMEA14430910 | 80 | 82 | 80 | 80 | 81 | 79 | 81 | 80 | 81 | 72 | 81 | 79 | 85 | 80 | 124 | 123 | 118 | 116 | 122 | 130 | 118 | 132 | 81 | 83 |

## References

1. 2023. Antimicrobial resistance in the EU/EEA (EARS-Net) - Annual Epidemiological Report 2022. European Centre for Disease Prevention and Control, Stockholm: ECDC.

2. Murray CJL, Ikuta KS, Sharara F, Swetschinski L, Robles Aguilar G, Gray A, Han C, Bisignano C, Rao P, Wool E, Johnson SC, Browne AJ, Chipeta MG, Fell F, Hackett S, Haines-Woodhouse G, Kashef Hamadani BH, Kumaran EAP, McManigal B, Achalapong S, Agarwal R, Akech S, Albertson S, Amuasi J, Andrews J, Aravkin A, Ashley E, Babin F-X, Bailey F, Baker S, Basnyat B, Bekker A, Bender R, Berkley JA, Bethou A, Bielicki J, Boonkasidecha S, Bukosia J, Carvalheiro C, Castañeda-Orjuela C, Chansamouth V, Chaurasia S, Chiurchiù S, Chowdhury F, Clotaire Donatien R, Cook AJ, Cooper B, Cressey TR, Criollo-Mora E, Cunningham M, Darboe S, Day NPJ, De Luca M, Dokova K, Dramowski A, Dunachie SJ, Duong Bich T, Eckmanns T, Eibach D, Emami A, Feasey N, Fisher-Pearson N, Forrest K, Garcia C, Garrett D, Gastmeier P, Giref AZ, Greer RC, Gupta V, Haller S, Haselbeck A, Hay SI, Holm M, Hopkins S, Hsia Y, Iregbu KC, Jacobs J, Jarovsky D, Javanmardi F, Jenney AWJ, Khorana M, Khusuwan S, Kissoon N, Kobeissi E, Kostyanev T, Krapp F, Krumkamp R, Kumar A, Kyu HH, Lim C, Lim K, Limmathurotsakul D, Loftus MJ, Lunn M, Ma J, Manoharan A, Marks F, May J, Mayxay M, Mturi N, Munera-Huertas T, Musicha P, Musila LA, Mussi-Pinhata MM, Naidu RN, Nakamura T, Nanavati R, Nangia S, Newton P, Ngoun C, Novotney A, Nwakanma D, Obiero CW, Ochoa TJ, Olivas-Martinez A, Olliaro P, Ooko E, Ortiz-Brizuela E, Ounchanum P, Pak GD, Paredes JL, Peleg AY, Perrone C, Phe T, Phommasone K, Plakkal N, Ponce-de-Leon A, Raad M, Ramdin T, Rattanavong S, Riddell A, Roberts T, Robotham JV, Roca A, Rosenthal VD, Rudd KE, Russell N, Sader HS, Saengchan W, Schnall J, Scott JAG, Seekaew S, Sharland M, Shivamallappa M, Sifuentes-Osornio J, Simpson AJ, Steenkeste N, Stewardson AJ, Stoeva T, Tasak N, Thaiprakong A, Thwaites G, Tigoi C, Turner C, Turner P, Van Doorn HR, Velaphi S, Vongpradith A, Vongsouvath M, Vu H, Walsh T, Walson JL, Waner S, Wangrangsimakul T, Wannapinij P, Wozniak T, Young Sharma TEMW, Yu KC, Zheng P, Sartorius B, Lopez AD, Stergachis A, Moore C, Dolecek C, Naghavi M. 2022. Global burden of bacterial antimicrobial resistance in 2019: a systematic analysis. The Lancet 399:629–655.

3. David S, Reuter S, Harris SR, Glasner C, Feltwell T, Argimon S, Abudahab K, Goater R, Giani T, Errico G, Aspbury M, Sjunnebo S, EuSCAPE Working Group, ESGEM Study Group, Feil EJ, Rossolini GM, Aanensen DM, Grundmann H. 2019. Epidemic of carbapenem-resistant Klebsiella pneumoniae in Europe is driven by nosocomial spread. Nat Microbiol 4:1919–1929.

4. Tsioutis C, Eichel VM, Mutters NT. 2021. Transmission of *Klebsiella pneumoniae* carbapenemase (KPC)-producing *Klebsiella pneumoniae* : the role of infection control. J Antimicrob Chemother 76:i4–i11.

5. Tacconelli E, Buhl M, Humphreys H, Malek V, Presterl E, Rodriguez-Baño J, Vos MC, Zingg W, Mutters NT. 2019. Analysis of the challenges in implementing guidelines to prevent the spread of multidrug-resistant gram-negatives in Europe. BMJ Open 9:e027683.

6. Baker KS, Jauneikaite E, Hopkins KL, Lo SW, Sánchez-Busó L, Getino M, Howden BP, Holt KE, Musila LA, Hendriksen RS, Amoako DG, Aanensen DM, Okeke IN, Egyir B, Nunn JG, Midega JT, Feasey NA, Peacock SJ. 2023. Genomics for public health and international surveillance of antimicrobial resistance. Lancet Microbe 4:e1047–e1055.

7. European Health and Digital Executive Agency., Tetra Tech. 2023. Study on the barriers to effective development and implementation of national policies on antimicrobial resistance: final report. Publications Office, LU. https://data.europa.eu/doi/10.2925/826400. Retrieved 14 May 2024.

8. Simar SR, Hanson BM, Arias CA. 2021. Techniques in bacterial strain typing: past, present, and future. Curr Opin Infect Dis 34:339–345.

9. Cameron A, Bohrhunter JL, Taffner S, Malek A, Pecora ND. 2020. Clinical Pathogen Genomics. Clin Lab Med 40:447–458.

10. Werner G, Couto N, Feil EJ, Novais A, Hegstad K, Howden BP, Friedrich AW, Reuter S. 2023. Taking hospital pathogen surveillance to the next level. Microb Genomics 9.

11. McGann P, Bunin JL, Snesrud E, Singh S, Maybank R, Ong AC, Kwak YI, Seronello S, Clifford RJ, Hinkle M, Yamada S, Barnhill J, Lesho E. 2016. Real time application of whole genome sequencing for outbreak investigation – What is an achievable turnaround time? Diagn Microbiol Infect Dis 85:277–282.

12. Price V, Ngwira LG, Lewis JM, Baker KS, Peacock SJ, Jauneikaite E, Feasey N. 2023. A systematic review of economic evaluations of whole-genome sequencing for the surveillance of bacterial pathogens. Microb Genomics 9.

13. Forde BM, Bergh H, Cuddihy T, Hajkowicz K, Hurst T, Playford EG, Henderson BC, Runnegar N, Clark J, Jennison AV, Moss S, Hume A, Leroux H, Beatson SA, Paterson DL, Harris PNA. 2023. Clinical Implementation of Routine Whole-genome Sequencing for Hospital Infection Control of Multi-drug Resistant Pathogens. Clin Infect Dis 76:e1277–e1284.

14. Foster-Nyarko E, Cottingham H, Wick RR, Judd LM, Lam MMC, Wyres KL, Stanton TD, Tsang KK, David S, Aanensen DM, Brisse S, Holt KE. 2023. Nanopore-only assemblies for genomic surveillance of the global priority drug-resistant pathogen, Klebsiella pneumoniae. Microb Genomics 9.

15. Wagner GE, Dabernig-Heinz J, Lipp M, Cabal A, Simantzik J, Kohl M, Scheiber M, Lichtenegger S, Ehricht R, Leitner E, Ruppitsch W, Steinmetz I. 2023. Real-Time Nanopore Q20+ Sequencing Enables Extremely Fast and Accurate Core Genome MLST Typing and Democratizes Access to High-Resolution Bacterial Pathogen Surveillance. J Clin Microbiol 61:e01631–22.

16. Novais Â, Freitas AR, Rodrigues C, Peixe L. 2019. Fourier transform infrared spectroscopy: unlocking fundamentals and prospects for bacterial strain typing. Eur J Clin Microbiol Infect Dis 38:427–448.

17. Martak D, Valot B, Sauget M, Cholley P, Thouverez M, Bertrand X, Hocquet D. 2019. Fourier-Transform InfraRed Spectroscopy Can Quickly Type Gram-Negative Bacilli Responsible for Hospital Outbreaks. Front Microbiol 10:1440.

18. Hu Y, Zhou H, Lu J, Sun Q, Liu C, Zeng Y, Zhang R. 2021. Evaluation of the IR Biotyper for *Klebsiella pneumoniae* typing and its potentials in hospital hygiene management. Microb Biotechnol 14:1343–1352.

19. Rodrigues C, Sousa C, Lopes JA, Novais Â, Peixe L. 2020. A Front Line on Klebsiella pneumoniae Capsular Polysaccharide Knowledge: Fourier Transform Infrared Spectroscopy as an Accurate and Fast Typing Tool. mSystems 5.

20. Novais Â, Gonçalves AB, Ribeiro TG, Freitas AR, Méndez G, Mancera L, Read A, Alves V, López-Cerero L, Rodríguez-Baño J, Pascual Á, Peixe L. 2024. Development and validation of a quick, automated, and reproducible ATR FT-IR spectroscopy machine-learning model for *Klebsiella pneumoniae* typing. J Clin Microbiol 62:e01211–23.

21. Silva L, Rodrigues C, Lira A, Leão M, Mota M, Lopes P, Novais Â, Peixe L. 2020. Fourier transform infrared (FT-IR) spectroscopy typing: a real-time analysis of an outbreak by carbapenem-resistant Klebsiella pneumoniae. Eur J Clin Microbiol Infect Dis 39:2471–2475.

22. Novais Â, Ferraz RV, Viana M, da Costa PM, Peixe L. 2022. NDM-1 Introduction in Portugal through a ST11 KL105 Klebsiella pneumoniae Widespread in Europe. Antibiot Basel Switz 11:92.

23. Brisse S, Passet V, Haugaard AB, Babosan A, Kassis-Chikhani N, Struve C, Decré D. 2013. *wzi* Gene Sequencing, a Rapid Method for Determination of Capsular Type for Klebsiella Strains. J Clin Microbiol 51:4073–4078.

24. Rodrigues C, Bavlovič J, Machado E, Amorim J, Peixe L, Novais Â. 2016. KPC-3-Producing Klebsiella pneumoniae in Portugal Linked to Previously Circulating Non-CG258 Lineages and Uncommon Genetic Platforms (Tn4401d-IncFIA and Tn4401d-IncN). Front Microbiol 7.

25. Henius A. 2023. EpiLinx_ECCMID2023_firstdraft.pdf. figshare.

26. Andrews S. 2010. FastQC: a quality control tool for high throughput sequence data.

27. Lam MMC, Wick RR, Judd LM, Holt KE, Wyres KL. 2022. Kaptive 2.0: updated capsule and lipopolysaccharide locus typing for the Klebsiella pneumoniae species complex. Microb Genomics 8.

28. Diancourt L, Passet V, Verhoef J, Grimont PAD, Brisse S. 2005. Multilocus Sequence Typing of *Klebsiella pneumoniae* Nosocomial Isolates. J Clin Microbiol 43:4178–4182.

29. Hennart M, Guglielmini J, Bridel S, Maiden MCJ, Jolley KA, Criscuolo A, Brisse S. 2022. A Dual Barcoding Approach to Bacterial Strain Nomenclature: Genomic Taxonomy of *Klebsiella pneumoniae* Strains. Mol Biol Evol 39:msac135.

30. Lam MMC, Wick RR, Watts SC, Cerdeira LT, Wyres KL, Holt KE. 2021. A genomic surveillance framework and genotyping tool for Klebsiella pneumoniae and its related species complex. Nat Commun 12:4188.

31. Carattoli A, Zankari E, García-Fernández A, Voldby Larsen M, Lund O, Villa L, Møller Aarestrup F, Hasman H. 2014. *In* *Silico* Detection and Typing of Plasmids using PlasmidFinder and Plasmid Multilocus Sequence Typing. Antimicrob Agents Chemother 58:3895–3903.

32. Argimón S, David S, Underwood A, Abrudan M, Wheeler NE, Kekre M, Abudahab K, Yeats CA, Goater R, Taylor B, Harste H, Muddyman D, Feil EJ, Brisse S, Holt K, Donado-Godoy P, Ravikumar KL, Okeke IN, Carlos C, Aanensen DM, NIHR Global Health Research Unit on Genomic Surveillance of Antimicrobial Resistance, Fabian Bernal J, Arevalo A, Fernanda Valencia M, Osma Castro ECD, Nagaraj G, Shamanna V, Govindan V, Prabhu A, Sravani D, Shincy MR, Rose S, Ravishankar KN, Oaikhena AO, Afolayan AO, Ajiboye JJ, Ewomazino Odih E, Lagrada ML, Macaranas PKV, Olorosa AM, Gayeta JM, Masim MAL, Herrera EM, Molloy A, Stelling J. 2021. Rapid Genomic Characterization and Global Surveillance of *Klebsiella* Using Pathogenwatch. Clin Infect Dis 73:S325–S335.

33. Yang H, Shi H, Feng B, Wang L, Chen L, Alvarez-Ordóñez A, Zhang L, Shen H, Zhu J, Yang S, Ding C, Prietod M, Yang F, Yu S. 2023. Protocol for bacterial typing using Fourier transform infrared spectroscopy. STAR Protoc 4:102223.

34. Ernst CM, Braxton JR, Rodriguez-Osorio CA, Zagieboylo AP, Li L, Pironti A, Manson AL, Nair AV, Benson M, Cummins K, Clatworthy AE, Earl AM, Cosimi LA, Hung DT. 2020. Adaptive evolution of virulence and persistence in carbapenem-resistant Klebsiella pneumoniae. Nat Med 26:705–711.

35. He J, Shi Q, Chen Z, Zhang W, Lan P, Xu Q, Hu H, Chen Q, Fan J, Jiang Y, Loh B, Leptihn S, Zou Q, Zhang J, Yu Y, Hua X. 2023. Opposite evolution of pathogenicity driven by in vivo wzc and wcaJ mutations in ST11-KL64 carbapenem-resistant Klebsiella pneumoniae. Drug Resist Updat 66:100891.

36. Guerra AM, Lira A, Lameirão A, Selaru A, Abreu G, Lopes P, Mota M, Novais Â, Peixe L. 2020. Multiplicity of Carbapenemase-Producers Three Years after a KPC-3-Producing K. pneumoniae ST147-K64 Hospital Outbreak. Antibiotics 9:806.

37. Spadar A, Phelan J, Elias R, Modesto A, Caneiras C, Marques C, Lito L, Pinto M, Cavaco-Silva P, Ferreira H, Pomba C, Da Silva GJ, Saavedra MJ, Melo-Cristino J, Duarte A, Campino S, Perdigão J, Clark TG. 2022. Genomic epidemiological analysis of Klebsiella pneumoniae from Portuguese hospitals reveals insights into circulating antimicrobial resistance. Sci Rep 12:13791.

38. Rodrigues C, Lanza VF, Peixe L, Coque TM, Novais Â. 2023. Phylogenomics of Globally Spread Clonal Groups 14 and 15 of Klebsiella pneumoniae. Microbiol Spectr 11:e03395–22.

39. Lin Y-T, Wang Y-P, Wang F-D, Fung C-P. 2015. Community-onset Klebsiella pneumoniae pneumonia in Taiwan: clinical features of the disease and associated microbiological characteristics of isolates from pneumonia and nasopharynx. Front Microbiol 9.

40. Zhang Y, Zhao C, Wang Q, Wang X, Chen H, Li H, Zhang F, Li S, Wang R, Wang H. 2016. High Prevalence of Hypervirulent Klebsiella pneumoniae Infection in China: Geographic Distribution, Clinical Characteristics, and Antimicrobial Resistance. Antimicrob Agents Chemother 60:6115–6120.

41. Faria NA, Touret T, Simões AS, Palos C, Bispo S, Cristino JM, Ramirez M, Carriço J, Pinto M, Toscano C, Gonçalves E, Gonçalves ML, Costa A, Araújo M, Duarte A, De Lencastre H, Serrano M, Sá-Leão R, Miragaia M. 2024. Genomic insights into the expansion of carbapenem-resistant Klebsiella pneumoniae within Portuguese hospitals. J Hosp Infect 148:62–76.

42. Martin RM, Cao J, Brisse S, Passet V, Wu W, Zhao L, Malani PN, Rao K, Bachman MA. 2016. Molecular Epidemiology of Colonizing and Infecting Isolates of Klebsiella pneumoniae. mSphere 1:e00261–16.

